# Combining fusion of cells with CRISPR-Cas9 editing for the cloning of large DNA fragments or complete bacterial genomes in yeast

**DOI:** 10.1101/2023.03.14.531922

**Authors:** Gabrielle Guesdon, Géraldine Gourgues, Fabien Rideau, Thomas Ipoutcha, Lucía Manso-Silván, Matthieu Jules, Pascal Sirand-Pugnet, Alain Blanchard, Carole Lartigue

## Abstract

The genetic engineering of genome fragments larger than 100 kbp is challenging and requires both specific methods and cloning hosts. The yeast *Saccharomyces cerevisiae* is considered as a host of choice for cloning and engineering whole or partial genomes from viruses, bacteria, and algae. Several methods are now available to perform these manipulations, each with its own limitations. In order to extend the range of in-yeast cloning strategies, a new approach combining two already described methods, the Fusion cloning and the CReasPy-Cloning, was developed. The CReasPy-Fusion method allows the simultaneous cloning and engineering of megabase-sized genomes in yeast by fusion of bacterial cells with yeast spheroplasts carrying the CRISPR-Cas9 system. With this new approach, we demonstrate the feasibility of cloning and editing whole genomes from several *Mycoplasma* species belonging to different phylogenetic groups. We also show that CReasPy-Fusion allows the capture of large genome fragments with high efficacy, resulting in the successful cloning of selected loci in yeast. We finally identify bacterial nuclease encoding genes as barriers for CReasPy-Fusion by showing that their removal from the donor genome improves cloning efficacy.

## Introduction

In Synthetic Biology, technologies are often developed using model organisms that are amenable to efficient genetic modifications, which acts as living workbenches. As such, the yeast *Saccharomyces cerevisiae* has long been used to propagate and edit genetic material from other organisms. In the late 80’s and for the first time, the cloning of linear DNA molecules with a size reaching 400 kbp as yeast artificial chromosomes (YACs)^1^, was achieved. This result opened up the possibility to clone large genome fragments from a wide range of organisms including eukaryotes, bacteria or viruses^2–4^. If this approach was a powerful step towards genome analysis, including physical maps of complex genomes^5, 6^, shotgun sequencing strategies^7^ or gene function studies^8^, instability issues of certain heterologous DNA fragments in yeast reduced its attractiveness and practical use.

It was not until the mid-2000s that yeast gained renewed interest as a cloning host. Thanks to its capacity to propagate longer DNA fragments more easily than other organisms such as *Escherichia coli* or *Bacillus subtilis*, the yeast *S. cerevisiae* was indeed chosen as the preferred cell host for the cloning of entire bacterial genomes^9, 10^ as circular yeast centromeric plasmids (YCps). A remarkable landmark was the complete assembly in yeast of the synthetic genome (582 kbp) of *Mycoplasma genitalium*, which perfectly illustrated these new possibilities. Using the efficient yeast homologous recombination machinery, the genome assembly was performed using 6 overlapping DNA fragments^11^ and reiterated a few months later with 25 overlapping fragments in a single step^12^. Following this work, many other partial or entire genomes (native or synthetic) from other bacteria, and even eukaryotes, were cloned into yeast as YCPs^10, 13–17^. Among them, the synthetic genomes *Mycoplasma mycoides* JCVI-syn1.0 and JCVI-syn3.0 which transplantation (from yeast to a recipient cell) resulted in the boot-up of the first synthetic cell^18^ and the first quasi-minimal synthetic cell, respectively^18, 19^.

More recently, the versatility of yeast as a host has been used for cloning and modifying viral genomes. This possibility was applied in the context of emerging viruses including the severe acute respiratory syndrome coronavirus 2 (SARS-CoV-2) that was achieved within only a few weeks after the release of SARS-CoV-2 genome sequence^20^. To date, there are no less than 25 bacterial and 10 viral genomes cloned in yeast^17, 21^.

The successful cloning of a whole genome in yeast requires to consider different key elements including (i) the characteristics of the donor genome organism, (ii) the donor genome itself (size, presence and number of restriction sites) and (iii) the downstream applications. In total, four different approaches are currently available^10, 12, 13, 22, 23^ ; all have in common the insertion of a yeast vector composed of a yeast centromere (CEN), a yeast selection marker and, in certain cases, a yeast origin of replication (ARS). In the conventional procedure, genomes marked with the yeast vector are isolated and transferred intact into yeast spheroplasts^10, 24^. In contrast, for the TAR-cloning^25–27^ and the CReasPy-Cloning protocols^23^, the yeast vector is directly transformed into yeast spheroplasts along with the bacterial genome. The common denominator of these three methods is the need to isolate intact naked genomes prior to yeast transformation. This critical step is performed in agarose plugs in order to protect DNA from shearing forces and avoid breakages as much as possible. Although this alleviates most of the problems, this step is still tedious. To get around this requirement, Karas *et al.* proposed the direct cell-to-cell transfer of genomes from bacteria to yeast spheroplasts. This protocol, initially developed using the wall-less bacteria belonging to the class *Mollicutes*, *Mycoplasma mycoides* subsp. *capri* (*Mmc*) and *Mycoplasma capricolum* subsp. *capricolum* (*Mcap*), was subsequently applied to the Gram-negative bacterium *Haemophilus influenzae*^22, 28^. Interestingly, this study showed that the removal of restriction endonucleases (RE) from the donor bacteria increased the fusion-mediated genome transfer efficacy. Later on, the same group revealed that genetic factors other than RE might also prevent the genome transfer from bacteria to yeast, since the deletion of the *glpF* gene (glycerol uptake facilitator protein) from the *M. mycoides* genome improved the efficacy of the method by up to 21-fold^29^. Although this method offers the advantage of not having to isolate genomes in agarose plugs, it has a major drawback: the yeast elements required for the maintenance and the replication of the bacterial genomes in yeast must be inserted in the bacterial genome before its transfer into yeast by cell fusion. This requirement strongly limits its broader application because transformation protocols are not always available for the bacteria whose genome is to be cloned.

In the present study, we extended the initial “Fusion Cloning” method of Karas *et al*. by combining the cell-to-cell transfer of genomes with the in-yeast “CReasPy-Cloning” method based on CRISPR-Cas9 and recently described^23^. By doing so, the insertion of the yeast elements occurs during the genome transfer from bacteria to yeast, eliminating the need for pre-marked bacterial genomes. The new method, named “CReasPy-Fusion”, consists of three main successive steps (**Figure 1**): (1) Yeast cells are transformed with a Cas9 expression plasmid (pCas9) and a gRNA expression plasmid (pgRNA) (step borrowed from CReasPy-Cloning); (2) Yeast cells harboring pCas9 and pgRNA are fused with the bacteria of interest (step borrowed from Fusion Cloning) in the presence of a specific recombination template; (3) After entry of the genome into yeast cells, the Cas9-gRNA duplex induces a double-strand break at the target site, which is repaired by the yeast homologous recombination system using the provided template. The repaired bacterial genome containing the yeast elements inserted at a precise locus, can now be propagated as a yeast centromeric plasmid. This method was validated using different *Mycoplasma* species of veterinary importance belonging to three distinct phylogenetic groups of the class *Mollicutes* (Spiroplasma, Pneumoniae and Hominis, **Figure S1**). Using this method, we also performed the cloning and concomitant inactivation of genes encoding recognized virulence factors for six out of seven species tested.

**Figure 1:**
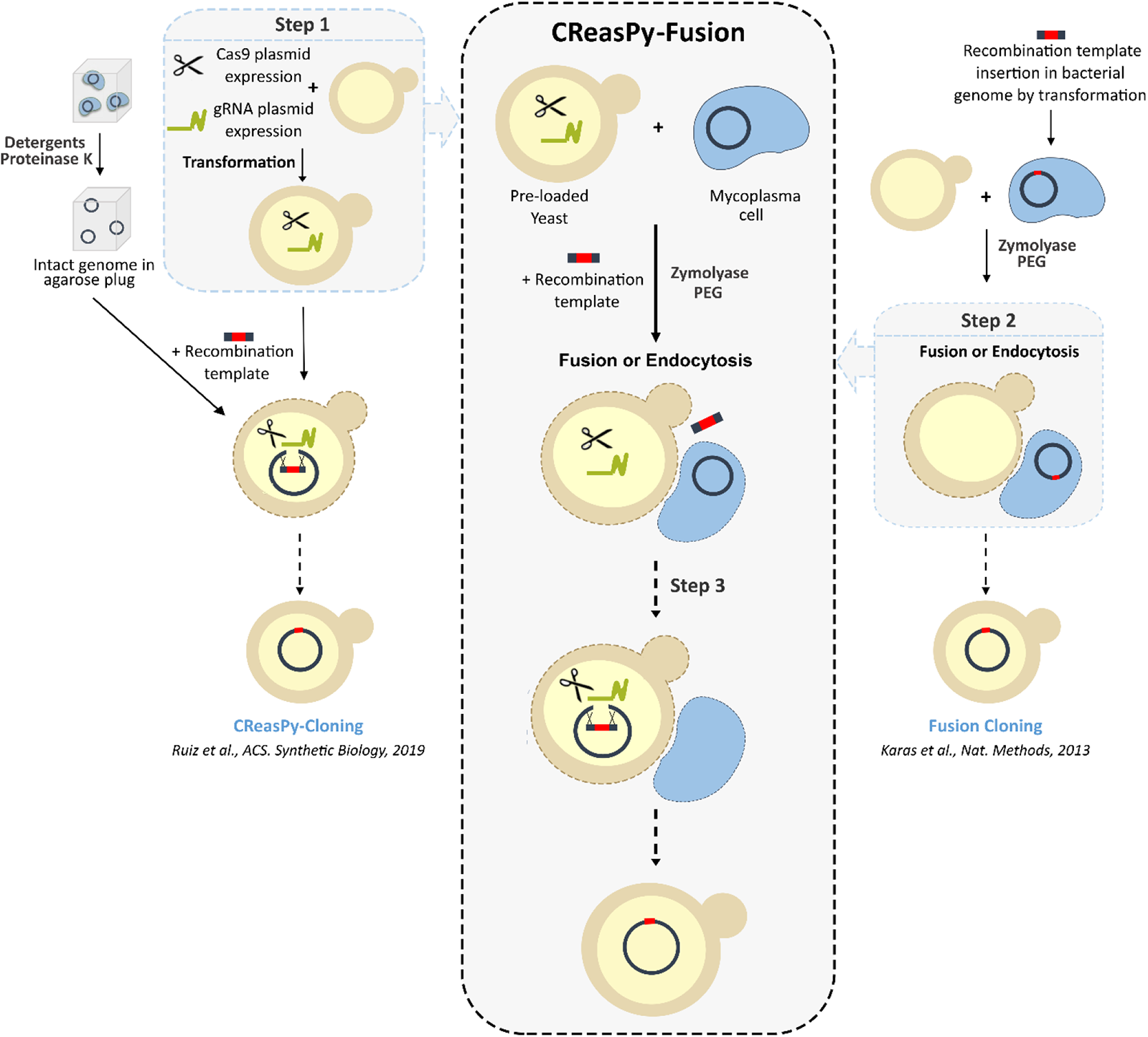
Schematic diagram of the experimental procedure of the CReasPy-Fusion method. Step 1 (borrowed from the CReasPy-Cloning strategy, left column): the yeast is transformed with two plasmids, allowing the expression of the Cas9 nuclease and a gRNA. Step 2 (borrowed from the Fusion Cloning strategy, right column): yeast cells preloaded with pCas9 and pgRNA are put in contact with mycoplasma cells in the presence of a linear recombination template (made of the yeast elements CEN-HIS3 with or without ARS flanked by two recombination arms identical to each side of the target locus and an antibiotic resistance marker). Step 3: Upon entry into the yeast cell, the target genome is cleaved by Cas9, and subsequently repaired by the yeast homologous recombination system using the provided linear DNA fragment as a template. As a result, the bacterial genome now includes the yeast elements inserted at a precise locus, and is carried by the yeast as a centromeric plasmid.

## Results and Discussion

CReasPy-Fusion experiments reported in this paper were performed using strains from seven *Mycoplasma* species (**Figure S1 and Table S1**), which can be divided into two groups: (i) species whose genomes had already been cloned in yeast using other procedures: *Mcap*, *Mmc*, and *M. mycoides* subsp. *mycoides* (*Mmm*)^23, 24^ and *M. capricolum* subsp. *capripneumoniae* (*Mccp*) (Personal communication from Dr. Carole Lartigue; Gourgues *et al.* unpublished), and (ii) species whose genomes had never been cloned in yeast before (*M. gallisepticum*, *M. agalactiae* and *M. bovis*).

### Simultaneous cloning and engineering of *Mycoplasma* genomes by CReasPy-Fusion: application to Mcap and Mmc, two mycoplasmas of the M. mycoides cluster

Before setting up the CReasPy-Fusion experiments, we wanted to make sure that we could reproduce the results previously published by Karas *et al.* in 2013^22^ and thus directly transfer whole marked genomes from bacteria to yeast by cell-to-cell fusion. We selected for this purpose the *Mcap* California Kid^T^ (CK^T^) strain mutant *Mcap*ΔRE^9^, in which the sole restriction system was inactivated by insertion in the encoding gene of the yeast elements and the puromycin resistance marker^30^. Following the authors’ instructions^22, 28^, yeast spheroplasts (strain VL6-48N) were co-incubated with *Mcap*ΔRE cells in the presence of polyethylene glycol (PEG) to promote cell fusion, then spread on selective yeast solid medium (SD-HIS). Depending on the conditions used, 88 to 202 yeast transformants were obtained (**Table S2**)^22, 28^. For each of four conditions tested, 10 colonies were randomly picked and analyzed by simplex PCR to detect the presence of the *McapΔRE* genome. Almost all clones (9/10 or 10/10) showed a band of 272 bp identical to that obtained with the positive control (*McapΔRE* gDNA) (**Figure S2**). Five clones per condition were then selected for multiplex PCR analysis, with the aim of verifying that the entire *McapΔRE* chromosome was potentially cloned in yeast. All transformants tested showed a 10-band profile (ranging from 100 bp to 1000 bp), identical to that of the positive control (**Figure S2**). Finally, two clones per condition were chosen to evaluate the size of the cloned DNA molecule by pulsed field gel electrophoresis (PFGE). All yeast clones displayed a two-band profile with sizes of 626 kbp and 383 kbp identical to that obtained with the *McapΔRE* positive control and corresponding to the theoretical sizes expected with the *McapΔRE* genome after restriction using the BssHII enzyme (**Figure S2**). In conclusion, almost all of the yeast transformants tested were found to correspond to yeast clones replicating the mycoplasma genome, thus validating the implementation of the protocol in the laboratory.

Based on these results, we attempted to merge the fusion method^22^ with the CReasPy-cloning method^23^ in order to develop the CReasPy-Fusion method (**Figure 1**), using *Mcap* and *Mmc*, two mycoplasmas belonging to the phylogenetic group Spiroplasma and, more specifically, to the *M. mycoides* cluster. For the first set of experiments, yeast spheroplasts preloaded with plasmids allowing the expression of the Cas9 nuclease and a gRNA were mixed with cells either from *Mcap* WT (strain CK^T^) or from *Mmc* WT (strain GM12) in the presence of a recombination template specific to the target site on the mycoplasma genome (**Figure 1**). For both species, the gene encoding the CCATC type II restriction endonuclease was chosen as target site for yeast vector and tetracycline resistance marker insertion (respectively MCAP_RS00270 and MMCAP2_RS00520). After transformation, the yeast transformants were screened as above, *i.e.*, first by simplex PCR to detect the presence of the genome and then by multiplex PCR and PFGE to check its integrity. All the results are shown in **Table 1** and an example of screening results is shown in **Figure S3**. Yeast carrying a complete mycoplasma genome were recovered both for *Mcap* and *Mmc* species, but with lower efficacies than those obtained during the Fusion experiments. For example, the number of colonies counted on the *Mcap* CK^T^ plate was ∼20 times lower with this method (10 to 12 transformants) than with the Fusion protocol (∼200 transformants). Among the 22 yeast transformants that were recovered, only 2 were shown to replicate the whole *Mcap* CK^T^ genome. Such a difference can probably be explained by the fact that during CReasPy-Fusion the genome must not only enter the host cell but also be modified to be stably maintained in the host. As observed for the CReasPy-Cloning method, to obtain yeast transformants propagating a mycoplasma genome a cascade of events must take place^23^: (i) transfer of the genome into yeast by fusion and concomitant acquisition of the recombination template, (ii) migration of both molecules into the yeast nucleus, (iii) double-stranded DNA cleavage of the genome by the Cas9-gRNA duplex, (iii) repair of the double-strand break by the yeast homologous recombination system using the provided template and (iv) maintenance and propagation of the bacterial genome (carrying the yeast elements inserted at a specific locus) as a centromeric yeast plasmid (**Table 1**).

**Table 1:** Screening of yeast transformants carrying Mcap and Mmc genomes generated by CReasPy-Fusion.

| *Mycoplasma* species | Experiments | CFU | Positive clones / Analyzed clones<sup>a</sup> |  |  |
| --- | --- | --- | --- | --- | --- |
|  |  |  | Simplex PCR | Multiplex PCR | PFGE<sup>b</sup> |
| *Mcap* (strain CK<sup>T</sup>) | Exp. 1 | 12 | 1/12 | 1/1 | 1/1 |
|  | Exp. 2 | 10 | 3/10 | 3/3 | 1/3 |
|  | TOTAL | 22 | 4/22 | 4/4 | 2/4 |
| *Mmc* (strain GM12) | Exp. 1 | 24 | 2/24 | 2/2 | 1/2 |
|  | Exp. 2 | 41 | 4/19 | 2/4 | 1/2 |
|  | TOTAL | 65 | 6/43 | 4/6 | 2/4 |

Two independent CReasPy-Fusion experiments were performed for each *Mycoplasma* species (*Mcap* or *Mmc*) as a replicate experiment. <sup>a</sup>The number of yeast transformants analyzed by simplex PCR, multiplex PCR and PFGE is reported, as well as the number of positive clones obtained. <sup>b</sup>PFGE analysis was not performed for all the positive clones but for a representative set of samples, as indicated in the table. *Mcap*: *M. capricolum* subsp. *capricolum*; *Mmc*: *M. mycoides* subsp. *capri*.

In order to demonstrate that mycoplasma genomes cloned using CReasPy-Fusion did not contain major mutations and were suitable for genome transplantation^9, 31^, edited *Mcap* and *Mmc* genomes were isolated from yeast clones, and subsequently transplanted in recipient *Mcap* cells (**Table S3**). A total of 3 and 12 putative bacterial transplants were obtained for *Mcap* and *Mmc* respectively and analyzed by species-specific multiplex PCR; all were identified as edited *Mcap* and *Mmc*. One transplant per species was selected for whole genome sequencing (cl 12.1 for *Mcap* and cl5.1 for *Mmc*). The analyses performed using Galaxy (https://usegalaxy.eu/) showed that sequences matched the expected genome design (without the targeted genes) and that no major recombination occurred after in-yeast CReasPy-Fusion editing. For both transplants, some SNPs and indels were identified (**Table S11**) but did not affect cell viability, consistent with the fact that most SNPs were silent mutations and probably related to natural error rate linked to host genome replication. Finally, these experiments confirmed that the CReasPy-Fusion method can be used for the production of mycoplasma mutant strains, in addition to other in-yeast cloning methods^9, 23, 32^.

### Extending the CReasPy-Fusion method to *Mmm* and *Mccp,* two major pathogens of the *M.* mycoides cluster

The results obtained with *Mcap* WT CK^T^ and *Mmc* WT GM12 led us to attempt an extension of the method to *Mccp* and *Mmm*^33^, two other mycoplasmas of the *M. mycoides* cluster of major veterinary importance (**Figure S1**). For *Mccp*, we selected two field strains^34^, the 95043 strain (isolated in Niger) and the 14020 strain (isolated in Tanzania). For *Mmm*, we selected the T1/44 vaccine strain^35^. Independent CReasPy-Fusion experiments were carried out for each strain and for each experiment the presence of the bacterial genome in yeast transformants was checked as above. The results obtained are summarized in **Table 2**.

**Table 2:**
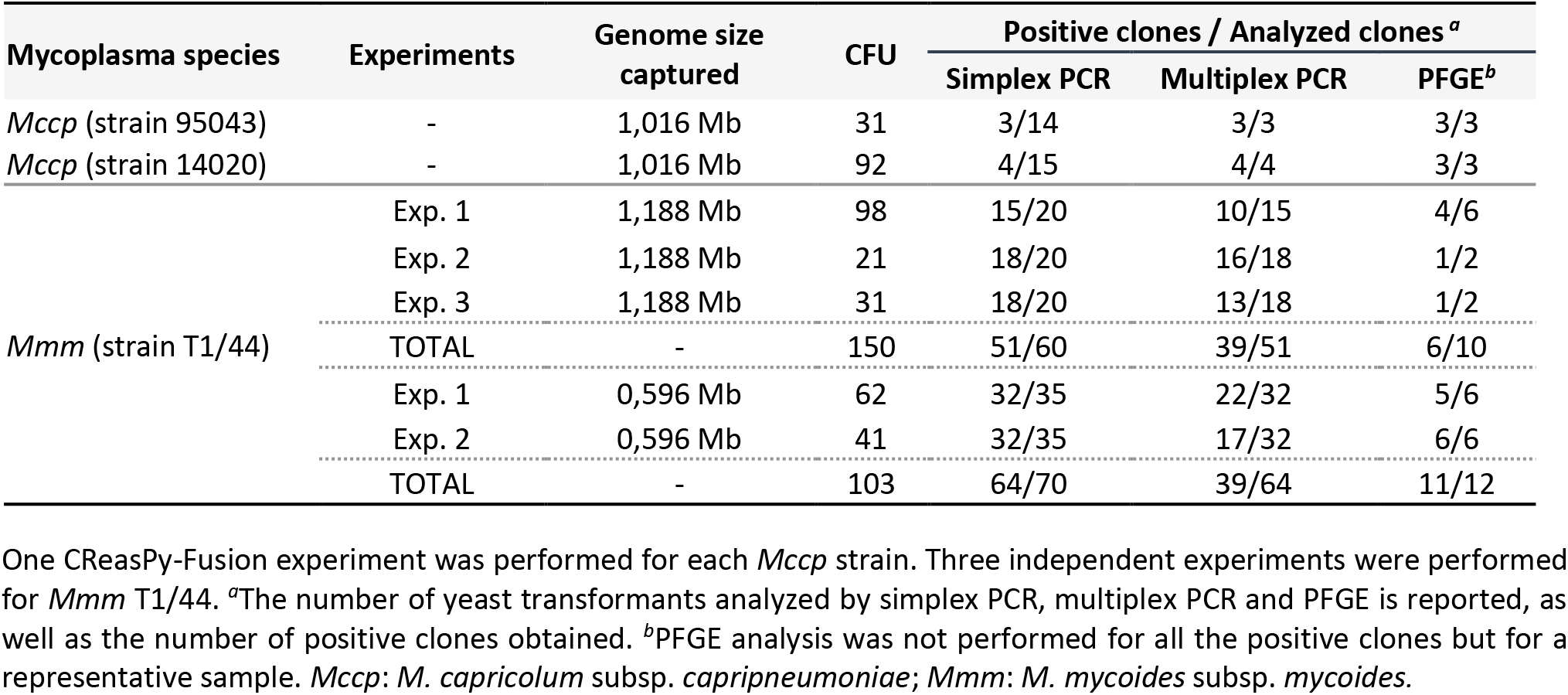
Screening of yeast transformants generated by CReasPy-Fusion using two major pathogenic mycoplasmas: Mccp and Mmm.

| Mycoplasma species | Experiments | Genome size captured | CFU | Positive clones / Analyzed clones <sup>a</sup> |  |  |
| --- | --- | --- | --- | --- | --- | --- |
|  |  |  |  | Simplex PCR | Multiplex PCR | PFGE<sup>b</sup> |
| *Mccp* (strain 95043) | - | 1,016 Mb | 31 | 3/14 | 3/3 | 3/3 |
| *Mccp* (strain 14020) | - | 1,016 Mb | 92 | 4/15 | 4/4 | 3/3 |
| *Mmm* (strain T1/44) | Exp. 1 | 1,188 Mb | 98 | 15/20 | 10/15 | 4/6 |
|  | Exp. 2 | 1,188 Mb | 21 | 18/20 | 16/18 | 1/2 |
|  | Exp. 3 | 1,188 Mb | 31 | 18/20 | 13/18 | 1/2 |
|  | TOTAL | - | 150 | 51/60 | 39/51 | 6/10 |
|  | Exp. 1 | 0,596 Mb | 62 | 32/35 | 22/32 | 5/6 |
|  | Exp. 2 | 0,596 Mb | 41 | 32/35 | 17/32 | 6/6 |
|  | TOTAL | - | 103 | 64/70 | 39/64 | 11/12 |

One CReasPy-Fusion experiment was performed for each *Mccp* strain. Three independent experiments were performed for *Mmm* T1/44. <sup>a</sup>The number of yeast transformants analyzed by simplex PCR, multiplex PCR and PFGE is reported, as well as the number of positive clones obtained. <sup>b</sup>PFGE analysis was not performed for all the positive clones but for a representative sample. *Mccp*: *M. capricolum* subsp. *capripneumoniae*; *Mmm*: *M. mycoides* subsp. *mycoides*.

CReasPy-Fusion experiments performed with *Mccp* WT strains were based on an experimental design aiming at cloning their whole genomes by targeting the peptidase S41 encoding gene (FOY67_01295 for the Nigerien strain and Mccp14020TZ_02950 for the Tanzanian strain, **Figure S3C**)^36^. Simplex PCR at the target locus revealed that 3 clones out of 14 (strain 95043) and 4 out of 15 (strain 14020) showed a band of the expected size (5044 bp) **(Figure S3C)**. Multiplex PCR then confirmed that no major genomic rearrangements had occurred in these clones, as the expected seven-band profile was visible on agarose gel for all of them (**Figure S3C)**. Finally, the 6 yeast transformants (3 for *Mccp* 95043 and 3 for *Mccp* 14020) selected for PFGE analysis showed genome profiles that were identical in size to that of the positive control (1016 kbp) after hydrolysis with the BssHII enzyme **(Figure S3C)**. These experiments showed that the genomes of both *Mccp* strains were successfully cloned in yeast by CReasPy-Fusion. The efficacy of cloning was very similar, with about 20% of the yeast clones analyzed propagating a whole *Mccp* genome for both strains. As previously for *Mcap* and *Mmc*, edited *Mccp* genomes from these strains were isolated from yeast clones, and subsequently transplanted in a recipient *Mcap* cell (**Table S3**) following a modified protocol developed in the Mollicutes Team (Personal communication from Dr. Carole Lartigue; Gourgues *et al.* unpublished). A single transplant of the Tanzanian strain was selected for whole genome sequencing (cl4.2) and analyzed. Very few mutations and indels were identified (**Table S11**) and no sequence rearrangements were detected. Those analyses confirm once again that the CReasPy-Fusion method does not affect the integrity of the cloned genomes.

In the case of *Mmm* strain T1/44, several CReasPy-Fusion experiments were performed not only to clone and simultaneously modify its whole genome using a single gRNA (1,188 kbp), but also to capture half of its genome using two gRNAs (596 kbp) (**Figure 2A**). For the whole genome cloning experiments (done in triplicate), we chose to target the *glpOKF* operon which includes genes implicated in hydrogen peroxide (H2O2) production ^37^. For the genome fragment capture experiments, which were performed twice, we built a plasmid encoding two distinct gRNAs (pgRNAs) to simultaneously target the *glpOKF* operon and either the MSCT144_RS01980 or the MSCT144_RS01995 target genes (encoding a lipoprotein and a hypothetical protein, respectively). The results are presented in **Table 2**. During the simplex PCR screening, a total of 51 clones out of 60 tested (whole genome) and 64 out of 70 tested (half genome) were positive (*i.e.* ∼80 and ∼90% positive clones respectively) (**Table 2 and** **Figure 2B**). Then, during the multiplex PCR screening, 39 clones out of 51 (whole genome) and 39 out of 64 tested (half genome) were validated (*i.e.* ∼60 and ∼75% of positive clones respectively) (**Table 2 and** **Figure 2C**). Finally, among the transformants selected for PFGE analysis, 6 clones out of 10 (whole genome) and 11 out of 12 tested (half genome) showed the expected profile (*i.e*., ∼60 and ∼90% positive clones respectively) (**Table 2 and** **Figure 2D**). More precisely, clones 8.1, 8.5, 15.3, 15.4 (exp. 1), 6.2 (exp. 2) and 6.5 (exp. 3) were shown to propagate the *Mmm* T1/44 whole genome (1,188 kbp), while all clones tested except clone 20.2 were shown to propagate the *Mmm* T1/44 half genome (596 kbp). Clone 20.2 displayed, in addition to the captured fragment, another band between 700 and 800 kbp.

**Figure 2:**
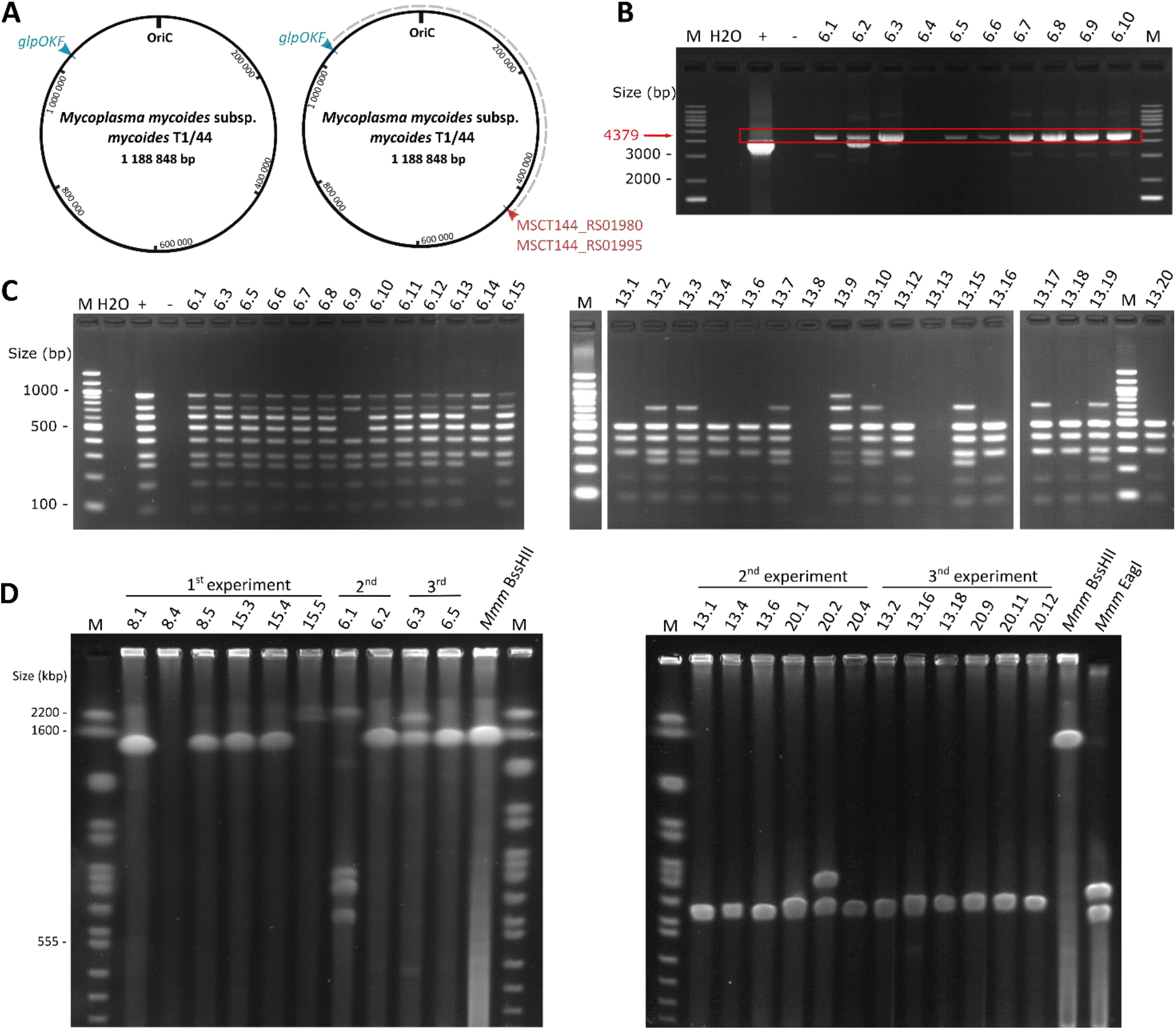
Screening of yeast transformants carrying either the entire *M. mycoides* subsp. *mycoides* (*Mmm*) genome or a genome fragment after CReasPy-Fusion. **(A)** Maps of *Mmm* T1/44 genome. The location of targeted loci (*glpOKF* operon, MSCT144_RS01980, MSCT144_RS01995) is indicated by coloured arrow heads. The grey dotted line indicates the half genome captured by CReasPy-Fusion. **(B)** The presence of the *Mmm* T1/44 genome in yeast and the expected target gene replacement were first assessed by simplex PCR analysis. The *glpOKF* operon deletion was validated by PCR using specific primers flanking the target locus. Positive transformants were validated with a 4,379 bp amplicon resulting from the insertion of the recombination template at the target site, instead of the 3,813 bp *glpOKF* amplicon expected for the WT strain. **(C)** Left panel: The completeness of the *Mmm* T1/44 genome cloned in yeast was assessed by multiplex PCR using a set of primers distributed around the genome (amplicons ranging from 89 bp to 1,020 bp). Clones carrying genomes without major rearrangement displayed a nine-band profile, identical to the one obtained for the positive control. Right panel: the capture of a half *Mmm* T1/44 genome in yeast was checked with the same multiplex primer set. Yeast transformants carrying the expected genome fragment (596 kbp) displayed a five-band profile (amplicons ranging from 89bp to 491 bp). **(D)** Left panel: the size of the entire *Mmm* T1/44 genome cloned in yeast was assessed by Pulsed Field Gel Electrophoresis (PFGE). Restriction of the bacterial genome with the enzyme BssHII should generate one linear DNA fragment of 1,200 kbp. Right panel: the size of the half *Mmm* T1/44 genome cloned in yeast was assessed in the same manner. Restriction of the bacterial genome with the enzyme BssHII should generate one linear DNA fragment of 596 kbp. For both experiments, positive controls consisting of *Mmm* T1/44 genomes isolated in agarose plugs and digested by either BssHII or EagI were loaded to the PFGE agarose gel. EagI digestion leads to two linear DNA fragments of 561 and 627 kbp, which are close to the size of the captured *Mmm* genome fragment (596 kbp). “M”: Simplex PCR = DNA Ladder 1kb+ Invitrogen (100-12,000 bp); Multiplex PCR: DNA Ladder 100bp NEB (100-1,517 bp); PFGE = *S. cervisae.* chromosomal DNA ladder Bio-Rad (225-2,200 kbp). “H2O”: negative control without DNA. “+”: positive control with *Mmm* T1/44 gDNA. “-“: negative control with *S. cerevisiae* VL6-48N gDNA.

The set of experiments conducted on *Mccp* and *Mmm* confirmed not only that (i) we were able to apply the CReasPy-Fusion method to clone genomes from field strains (*Mccp*) and from a vaccine strain (*Mmm*), but that (ii) we could also capture a genome fragment using two gRNAs. We might have thought that the efficacy of capturing a genome fragment would be lower, as is the case in CReasPy-Cloning, when several loci are targeted at the same time^23^. This was not the case at least in this example; indeed, the results obtained indicated that it was possible to circularize a portion of the chromosome after cutting two loci 500 kbp apart with an efficacy equivalent to that of cloning a whole genome after a single Cas9-mediated cleavage. Moreover, working with four species of the *M. mycoides* cluster (*Mccp/Mmm* here and *Mcap/Mmc* previously), we realized that cloning efficacies were highly variable from one species to another. Indeed, CReasPy-Fusion efficacies obtained with *Mccp* and *Mmm* species were remarkably high, while those obtained with *Mcap* and *Mmc* were rather low. We believe that this variation is due, in part, to the gRNA selected and consequently to the target gene or region. Indeed, gRNA cloned into pgRNA plasmids are selected to specifically target a chosen locus using Benchling (https://www.benchling.com/). This tool lists all 20-nucleotide sequences (or “spacers”) present upstream of a PAM (NGG) sequence in that given locus and ranks them using scores. These scores reflect the presumed efficacy of the gRNA during gene deletion experiments using the CRISPR-Cas9 tool. Sequences with ON target (spacer efficiency)^38^ and OFF target (spacer specificity)^39^ scores close to 100 are supposed to be the most efficient. Our long-term experience shows that gRNAs with high scores are sometimes completely inefficient *in vivo*, it is therefore very likely that among the gRNAs selected for our study, some are more effective than others^23^. Apart from the selection of the gRNA, we also believe that some of the parameters of the CReasPy-Fusion method may be better adjusted in order to optimize the protocol to each particular species. We can cite among other parameters: the bacterial growth phase, the nature and composition of the buffers used, and the bacterial protoplasts/yeast spheroplasts ratio. Some of these are key elements in the development of microorganism’s transformation protocols^40^; they are certainly key here as well. Adjusting these parameters to each particular species and strain may surely improve cell-to-cell fusion and, thus, the efficacy of the method.

### Overcoming a potential barrier for *M. gallisepticum* genome cloning using CReasPy-Fusion

In order to demonstrate the power and versatility of the CReasPy-Fusion method, we attempted an extension of its use by cloning and editing the genome of *M. gallisepticum* (strain S6^T^), a poultry pathogen belonging to the Pneumoniae phylogenetic group (**Figure S1**). Unlike the four previous species of the *M. mycoides* cluster, the cloning of the *M. gallisepticum* genome in yeast had not been previously described. We designed a gRNA for cloning its entire genome, while targeting the *cysP* gene encoding a protease (GCW_RS001695, **Figure 3A**) that has a specificity for chicken antibodies^41^ and is therefore considered as a putative virulence factor. Our first experiment using this approach remained unsuccessful (**Table 3 and Table S4**). We hypothesized that this failure may be due to particular barriers inhibiting the cloning of the *M. gallisepticum* genome in yeast.

**Figure 3:**
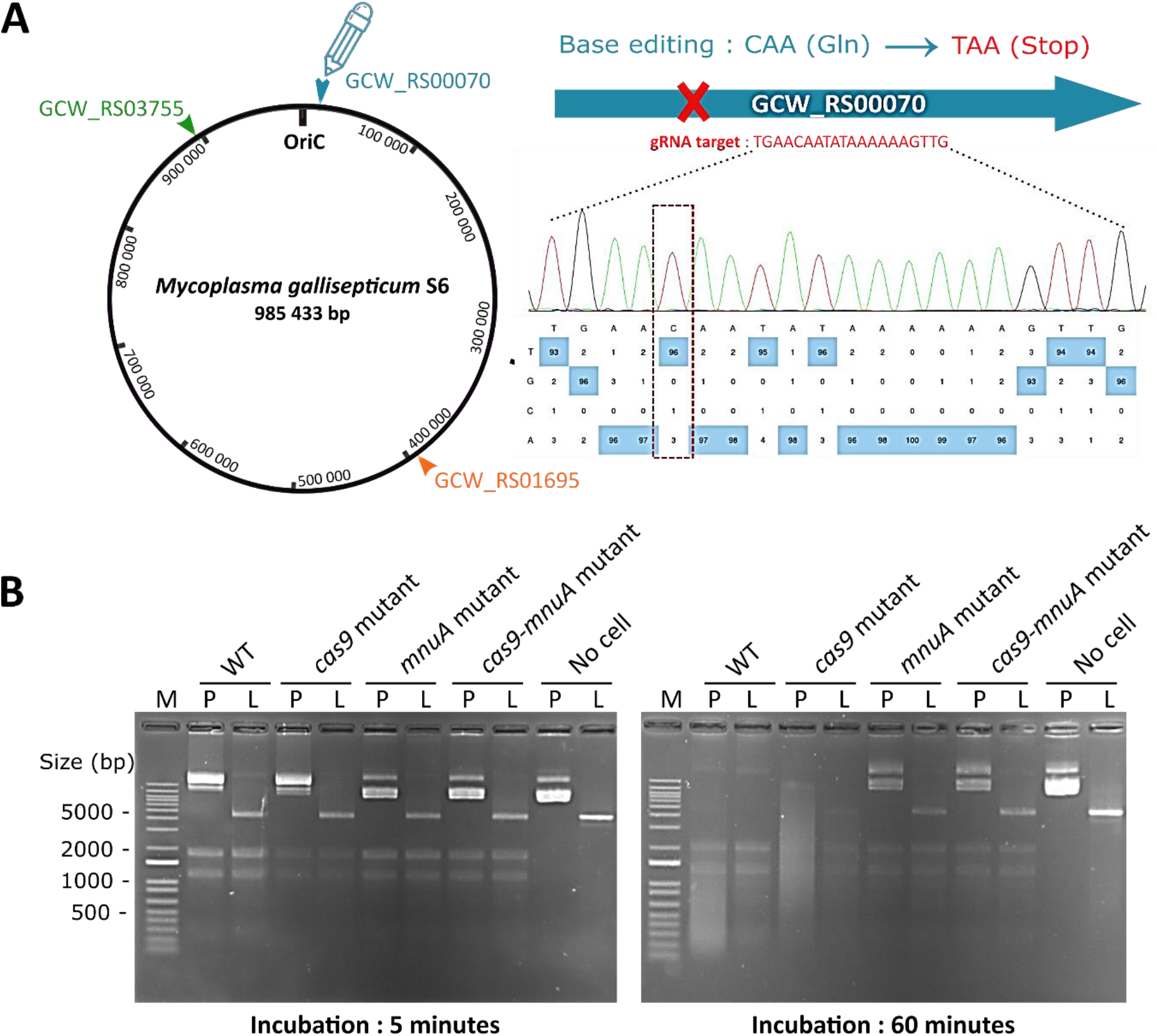
Inactivation of the nuclease membrane MnuA homolog in *M. gallisepticum* S6^T^ genome. **(A)** Map of *M. gallisepticum* S6^T^ genome. The location of targeted *loci* (GCW_RS00070 and GCW_RS01695) is indicated by coloured arrow heads (blue and orange respectively). The location of the Cas9 gene (GCW_RS03755) is indicated by a green arrow. Base editing of the *mnuA* gene (GCW_RS00070) is indicated by a blue pencil scheme: the gRNA was designed to replace a CAA glycine codon by a UAA stop codon resulting in the inactivation of the *mnuA* target gene. Chromatogram results associated with the EditR software analysis (https://moriaritylab.shinyapps.io/editr_v10/) are represented and the“C“ to “T” peak change is indicated by a rectangle dotted line **(B)** Nuclease activity test is shown for both wild-type and *M. gallisepticum* mutants at 5 minutes (left) and 60 minutes (right) of incubation. “M”: DNA Ladder 1kb+ Invitrogen (100-12000 bp); “P”: plasmid DNA; “L”: linear DNA.

**Table 3:** Screening of the yeast transformants generated after four independent CReasPy-Fusion experiments with *M. gallisepticum* WT strain and mutants.

| *Mycoplasma gallisepticum* donor genome | CFU | Positive clones / Analyzed clones<sup>a</sup> |  |  |
| --- | --- | --- | --- | --- |
|  |  | Simplex PCR | Multiplex PCR | PFGE<sup>b</sup> |
| 1<sup>st</sup> experiment |  |  |  |  |
| *M. gallisepticum* WT | 256 | 0/108 | - | - |
| 2<sup>nd</sup> experiment |  |  |  |  |
| *M. gallisepticum cas9-mnuA* mutant | 1036 | 12/117 | 11/12 | 2/2 |
| 3<sup>rd</sup> experiment |  |  |  |  |
| *M. gallisepticum* WT | 134 | 0/25 | - | - |
| *M. gallisepticum cas9* mutant | 82 | 0/27 | - | - |
| *M. gallisepticum mnuA* mutant | 115 | 3/34 | 2/3 | 1/1 |
| *M. gallisepticum cas9-mnuA* mutant | 84 | 7/27 | 4/7 | 1/1 |
| 4<sup>th</sup> experiment |  |  |  |  |
| *M. gallisepticum* WT | 298 | 1/16 | 1/1 | 1/1 |
| *M. gallisepticum cas9* mutant | 412 | 0/20 | - | - |
| *M. gallisepticum mnuA* mutant | 416 | 1/15 | 1/1 | 1/1 |
| *M. gallisepticum cas9-mnuA* mutant | 292 | 7/15 | 6/7 | 1/1 |

<sup>a</sup>The number of yeast transformants analyzed by simplex PCR, multiplex PCR and PFGE is reported, as well as the number of positive clones obtained. <sup>b</sup>PFGE analysis was not performed for all the positive clones but only for a representative sample.

Among the known cloning barriers, restriction modification (RM) systems are already known to lower the cell-to-cell genome transfer efficacy^22^. Using an analysis based on the Rebase Database^42^, two RM systems were identified **(Table S5).** However, similar RM systems are also found in the *Mmc, Mcap*, *Mmm* and *Mccp* genomes and they do not seem to inhibit cloning of the genome of these mycoplasmas. Therefore, we suspected that other putative barriers could be involved in this failure, including cytoplasmic, secreted or membrane nucleases with lower specificity than those associated to the RM systems. Indeed, we hypothesized that the specific recombination template used in our approach, provided as a linear double-stranded PCR product and carrying yeast elements, may be degraded by such nucleases. A search for nucleases other than those involved in RM systems yielded three candidates. The first one was the MnuA homolog, encoded by the CDS GCW_RS00070 locus. MnuA is a membrane nuclease characterized as a virulence factor in *M. bovis* (MBOVPG45_0215)^43^. MnuA is capable of digesting linear and circular DNA and requires activation by divalent cations (cofactors) such as Ca^2+^ and Mg^2+^, which are present in significant amounts in the buffers and solutions used here. This gene is absent in the genome of *M. mycoides* cluster species. The second candidate was the GCW_RS00180 protein identified as a Ca^2+^-dependent cytotoxic nuclease of *M. gallisepticum* with a staphylococcal nuclease region that displays the hallmarks of nucleases^44, 45^. The third candidate was the Cas9 endonuclease encoded by *M. gallisepticum* (GCW_RS03755) itself, which has been shown to be functional in previous studies^46, 47^.

In order to evaluate the possibility that nucleases inhibit genome cloning experiments, three mutants were selected to test the CReasPy-Fusion method. The first mutant was a spontaneous Cas9 deficient mutant previously obtained in the laboratory. In this mutant, a frameshift at the beginning of the *cas9* nucleotide sequence (position 51) leads to premature termination of the protein expression and loss of function^48^. The two other mutants (an *mnuA* mutant and a *cas9-mnuA* double mutant) (**Figure 3A**) were generated for this study using the CRISPR-Cas9 base editor system (CBE), which was recently adapted to mycoplasmas^49^. We chose not to include a “GCW_RS00180 mutant” initially because a study performed on *M. bovis* PG45^T^ had shown that among all nuclease coding genes in this species (3 in total), *mnuA* was responsible for the majority of the nuclease activity detectable *in vitro*^43^. We characterized the nuclease activity of the three mutants using an approach previously described for MnuA assay in *M. bovis* cells^43^. The effect of inactivating these nucleases was evaluated by co-incubating cells from both the WT and the mutants with either plasmid or linear DNA (**Figure 3B**). After 5 min of co-incubation there was no significant difference between the WT cells and those from the three mutants. However, following 1 hour of co-incubation, there was a striking difference with significantly higher DNA degradation observed in the WT and *cas9* mutant. Consistent with results obtained previously with *M. bovis*, the mutants affecting the membrane nuclease MnuA were those with the lower degree of DNA hydrolysis (**Figure 3B**).

We conducted a second CReasPy-Fusion experiment solely with the *cas9-mnuA* double mutant (**Table 3**). This choice was motivated by the fact that we wanted to get rid of both nucleases before determining whether one was more problematic than the other. During this experiment, we also evaluated if the protocol could be improved by: (i) addition of EDTA in the resuspension buffer of *M. gallisepticum* cells with the aim of chelating the divalent cations that are known cofactors of enzymes such as GCW_RS00180^45^; (ii) addition of single-stranded “cargo” DNA, such as denatured salmon sperm DNA in order to protect the repair template from nucleases by competition; and (iii) heating mycoplasma cells at 49°C to potentially inhibit some enzymatic activities^28^. The results obtained for each condition are presented in **Table S6**. A total of 1,036 colonies were obtained. When possible, ten colonies per condition were picked for further characterization. Considering that some of them did not regrow, we analyzed a total of 117 yeast colonies. After analysis by simplex PCR and multiplex PCR, we identified 11 transformants out of the 117 potentially propagating an entire genome. The two clones selected for PFGE analysis harbored a whole *M. gallisepticum* genome (clones 6.9 and 10.5; **Figure S4D**). Altogether these results show that nucleases constitute a barrier to the installation of the *M. gallisepticum* genome in yeast, either during or after cell-to-cell fusion. It can be stressed that none of the tested conditions resulted in clear improvement of the CReasPy-Fusion method over the previously defined protocol. These were therefore not retained for further experiments.

A third and a fourth experiment were performed by including, in addition to the *cas9-mnuA* double mutant, the *M. gallisepticum* WT strain as well as the *cas9* and *mnuA* single mutants (**Table 3**). In the third experiment, we successfully cloned the *M. gallisepticum* genome from both the *mnuA* mutant and the *cas9-mnuA* double mutant. For these two conditions, we obtained respectively: (i) 3 positive clones out of 34, and, 7 out of 27 in simplex PCR, then (ii) 2 clones out of 3, and, 4 out of 7 in multiplex PCR (**Table 3**). Finally, the two selected clones (one for each condition, *i.e.*, clones 16.3 and 20.8) analyzed by PFGE were validated, with a two-band profile of the expected size obtained after SacII digestion (536 kbp and 449 kbp; **Figure S4D**). In contrast, none of the clones tested for the *M. gallisepticum* WT strain and the *cas9* mutant were positive. This result may indicate that the enzyme preventing cloning of the *M. gallisepticum* genome in yeast is the surface nuclease MnuA, rather than the Cas9 protein. However, the fourth experiment forced us to moderate this conclusion. Indeed, we obtained positive clones for the *mnuA* mutant and the *cas9-mnuA* double mutant as before, but also for the WT strain of *M. gallisepticum* for which 1 (clone 9.3) out of 16 clones analyzed was validated by PFGE (**Table 3 and Figure S4**). On the other hand, only 1 (clone 13.1) out of 15 analyzed passed the three-steps screening and was validated for the *mnuA* mutant. Therefore, at this stage, we were not able to conclude whether the deletion of the MnuA nuclease-encoding gene completely unlocked cloning of the *M. gallisepticum* genome in yeast. However, the inactivation of *mnuA* combined with that of the *cas9* always led the recovery of a higher number of yeast transformants propagating the whole *M. gallisepticum* genome. The inactivation of the third nuclease (GCW_RS00180) may possibly result in increased efficacy. Altogether, our results confirmed that the CReasPy-Fusion method can be extended to mycoplasmas from phylogenetic groups other than Spiroplasma. This is indeed the first description of the cloning of the *M. gallisepticum* genome in yeast as a supernumerary centromeric plasmid.

### Assessment of the CReasPy-Fusion’s limitations using *Mycoplasma* species belonging to the Hominis phylogenetic group

Following the development of the CReasPy-Fusion method and the demonstration that membrane nucleases may act as inhibitors, we sought to extend the approach to two other *Mycoplasma* species whose genomes had not yet been cloned in yeast and which belong to a third phylogenetic group, Hominis (**Figure S1**). With this goal, we chose *M. bovis* PG45^T^ and *M. agalactiae* PG2^T^, two other mycoplasmas pathogenic for ruminants and that are phylogenetically close to each other. Taking into account the results obtained above, *mnuA* mutants of these mycoplasmas were produced by targeted base editing using Cas9 deaminase^49^. The gene encoding MnuA in *M. bovis* is MBOVPG45_0215 and its homolog in *M. agalactiae* is MAG_5900. While the phenotype of the *M. bovis mnuA* mutant was described in Ipoutcha *et al.*, 2022^49^, the phenotype of the *M. agalactiae mnuA* mutant is presented here (**Figure S5**). Just like *M. bovis* (and *M. gallisepticum* above), *M. agalactiae mnuA* inactivation resulted in a significant reduction of DNA hydrolysis when mutant cells were incubated with DNA substrates, either linear or circular.

Prior to the CReasPy-Fusion experiments, specific sgRNAs required to guide Cas9 cleavage in yeast were generated for these two species. The MBOVPG45_0215 and MAG_5900 *loci* encoding the MnuA nuclease were targeted in *M. bovis* PG45^T^ and *M. agalactiae* PG2^T^ respectively in order to completely remove these base edited genes at the time of the cloning. Two independent experiments were performed for each species. The results are summarized in **Table 4**. All the attempts for cloning the *M. bovis* genome remained unsuccessful. Indeed, all clones analyzed by simplex PCR remained negative, whether they were from *M. bovis* WT or *M. bovis mnuA* mutant (**Table S8 and S9**). As for *M. gallisepticum*, we attempted to counteract other potential harmful enzymatic activities using EDTA, salmon sperm DNA, trypsin limited treatment, but all experiments failed. It is possible that *M. bovis*/yeast fusion is less efficient than with other *Mycoplasma* species but, in our opinion, this does not seem to be the main reason for this failure. Indeed, *M. bovis* PG45^T^ possesses a very high number of RM systems. Karas *et al*. demonstrated that the presence of these systems could strongly decrease the efficiency of cell fusion^22^. In **Table S5**, we have listed all type I, II, and III RM systems predicted for all the species used during this work. We identified them using the online tool Rebase DataBase^42^ (http://rebase.neb.com/rebase/rebase.html), as well as the Molligen 3.0 and Molligen 4.0 databases (https://services.cbib.u-bordeaux.fr/molligen/ and http://www.molligen.org)^50^, and Padloc^51^ (https://padloc.otago.ac.nz/padloc/). In this table, we have specified whether the restrictases were described in the literature or identified as putative (on Molligen). For *M. bovis* PG45^T^, no less than 10 RM systems were predicted by Rebase and 9 restrictases (some of them referenced as truncated, *e.g.,* MBOVPG45_0615 and MBOVPG45_0617). It would be interesting to delete several of these loci in order to reduce the number of RM systems in *M. bovis* PG45^T^, and test the resulting mutants during a CReasPy-Fusion experiment. A simpler but less demonstrative strategy would be to identify an *M. bovis* strain that naturally has a reduced number of RM systems such as the strain RM16 with 5 type II systems. Apart from RM systems, other factors might also be involved, as discussed below.

**Table 4:** Comparison of positive yeast transformants obtained by CReasPy-Fusion using M. bovis WT and M. agalactiae WT and cognate mnuA mutants.

| *Mycoplasma* species | Experiments | CFU | Positive clones / Analyzed clones<sup>a</sup> |  |  |
| --- | --- | --- | --- | --- | --- |
|  |  |  | Simplex PCR | Multiplex PCR | PFGE |
| *M. bovis* (strain PG45<sup>T</sup> WT) | Exp. 1 | 740 | 0/92 | - | - |
| *M. bovis* (strain PG45<sup>T</sup> *mnuA* mutant) | Exp. 2 | 1414 | 0/320 | - | - |
| *M. agalactiae* (strain PG2<sup>T</sup> WT) | Same exp. | 523 | 2/43 | 2/2 | 2/2 |
| *M. agalactiae* (strain PG2<sup>T</sup> *mnuA* mutant) |  | 864 | 5/80 | 3/5 | 2/3 |

Two CReasPy-Fusion experiments were performed for *M. bovis* comparing both WT and *mnuA* mutant. One experiment was performed with *M. agalactiae* comparing both WT and *mnuA* mutant. <sup>a</sup>The number of yeast transformants analyzed by simplex PCR, multiplex PCR and PFGE is reported, as well as the number of positive clones obtained.

In contrast to *M. bovis*, the *M. agalactiae* genome was cloned using both the cells from WT and *mnuA* mutant strains (**Table 4 and Figure S10)**. No clear difference was observed between the two, suggesting that in contrary to *M. gallisepticum*, the activity of *M. agalactiae* MnuA does not interfere with the cloning process. In mycoplasmas, the first nuclease was identified approximately 20 years ago. Since then, a number of similar enzymes or homologous genes have also been reported^45^. Those characterized nucleases have been shown to possess different biological properties in terms of cofactors, substrates, inhibiting agents, subcellular location, etc. It is therefore possible that not all (secreted or membrane) nucleases be active under the conditions of this experiment, which can explain the differences observed in terms of cloning efficacy between the different *Mycoplasma* species handled during this study. In particular, it should be stressed out that amino acid sequence alignment of *M. bovis* and *M. agalactiae* MnuA proteins indicated that they are well conserved (75% identity) but differ from that of *M. gallisepticum* (30% identity). In addition, it is possible that these enzymes be more or less expressed or active depending on the growth phase or other conditions. All this may be sufficient to explain the difference observed between *M. gallisepticum* and *M. agalactiae* in CReasPy-Fusion.

Besides MnuA, other nucleases (*e.g.* GCW_RS00180 in *M. gallisepticum*, MAG_5040 in *M. agalactiae*, MBOVPG45_0310 or MBOVPG45_0089 in *M. bovis*) and unidentified factors may also impact CReasPy-Fusion. In a recent publication, Karas and co-authors, described that inactivation of the *glf* gene (encoding a protein involved in glycerol import), significantly increased genome transfer by Fusion^29^. This gene is present in all mycoplasma genomes manipulated in this study. This result remains poorly understood but suggests that other unknown factors may well play a role in the efficacy of CReasPy-Fusion. Membrane proteins may also, by their presence or absence, promote membrane-to-membrane fusion for example.

To conclude this section, even though no yeast transformants were obtained with the *M. bovis* genome, we succeeded in cloning the *M. agalactiae* genome for the first time in yeast.

## Conclusion

Over the past decade, yeast has regained a remarkable leading position for the cloning and editing of native or synthetically assembled genomes^17, 21^. The choice of the cloning method is essential and depends mainly on the micro-organism of interest and the characteristics of its genome. In order to increase the range of in-yeast cloning strategies, we developed here a new approach, named CReasPy-Fusion, that allows the simultaneous cloning and engineering of mega-sized genomes in yeast using the CRISPR-Cas9 system by direct bacterial cell to yeast spheroplast fusion.

For the development of this method, we chose to work with mycoplasmas. This choice was motivated by (i) the lack of a cell-wall in these bacteria, which should facilitate the fusion step with the yeast host cells, (ii) the small size of their genomes (<1.5 Mb), their low G+C content (≤40%), and the use of a non-standard genetic code (characteristics recognized as positive factors for the maintenance of genomes in *S. cerevisiae*^17, 52^). In addition, several mycoplasma genomes that had been previously cloned in yeast using different approaches including the Fusion method^22^ served as controls for the present study. Indeed, we knew that genomes from at least four out the seven species selected (*Mcap*, *Mmc*, *Mccp* and *Mmm*) could be cloned in yeast, and that the absence of transformants for these four species should have been attributed to a technical rather than a biological problem.

This work has allowed us to progress in the achievement and understanding of genome cloning procedures in yeast. First, the genome of six distinct *Mycoplasma* species, corresponding to seven strains distributed in three distinct phylogenetic groups were cloned/edited in yeast: *Mcap*, *Mmc*, *Mmm*, *Mccp* belonging to the Spiroplasma phylogenetic group, *M. gallisepticum* from the Pneumoniae group, and *M. agalactiae* from the Hominis group. For two of them, *M. gallisepticum* and *M. agalactiae*, this constitutes the first description of their genome cloning in yeast. Concerning *Mmm* and *Mccp*, the cloning of the genomes of *Mmm* strain PG1^T24^ and *Mccp* strain Abomsa (Personal communication from Dr. Carole Lartigue; Gourgues *et al.* unpublished) had already been described and the versatility of in-yeast cloning in these species is demonstrated here by further cloning of those of *Mmm* strain T1/44 and *Mccp* strains 14020 and 95043. Second, a derivative CReasPy-Fusion method has been developed allowing the capture of large genome fragments. This may be of particular interest to capture either part of more complex genome^53^, or large metabolic pathways. Third, we have demonstrated that, for the species *M. gallisepticum*, the inactivation of nuclease-encoding genes increased the number of yeast transformants, pointing out the impact of such enzymes in cell-to-cell genome transfer efficiency. Through our experiments, we realized that cloning efficacies were extremely variable between species, and even strains. These variations are most probably multifactorial: gRNA selection, species-specific factors, protocol settings… In this respect, Karas *et al.*, emphasize, for instance, the need to use early-exponential phase mycoplasma cultures to increase the frequency of genome transfer^28^.

We encountered difficulties in cloning and editing the genome of only one strain, *M. bovis* PG45^T^, which might be related to the high number of RM systems found in it. Indeed, while *Mcap* CK^T^ and *Mmc* GM12 genomes can be cloned both using CReasPy-Cloning (a method that uses genomes isolated in agarose plugs and deproteinized) and CReasPy-Fusion, the genomes of their counterparts *Mcap* 14232^54^ and *Mmc* 95010^55^, which have a different and more complex set of RM systems could not be cloned using CReasPy-Fusion despite several attempts (Lartigue *et al.*, personal communication). These results suggest that factors present in the cytoplasm, such as restrictases of RM systems and other less-specific nucleases, interfere with in-yeast genome cloning. In the case of *M. bovis*, as compared to *M. gallisepticum* and *M. agalactiae*, the use of the *mnuA* mutant made no difference, suggesting the presence of other cloning barriers.

Finally, with this work, we provide a new in-yeast cloning method that offers specific advantages over those previously published. In particular, the CReasPy-Fusion method can be used with bacteria that are not amenable to transformation or genetic modification. In addition, this approach alleviates the tedious steps of chromosome purification in agarose plugs which results in lowering both the time of preparation and the costs. Furthermore, since Karas *et al.*, showed that the genome of Gram-*Haemophilus influenzae* can be cloned in yeast by cell-to-cell Fusion, it would be of interest to expand CReasPy-Fusion to other Gram- and Gram+ bacteria. It is possible that for large genomes (>2 Mb), the Fusion method and CReasPy-Fusion are the most appropriate.

## Methods

### Yeast and Bacterial Strains, culture conditions

The *Mycoplasma* species and strains used in this study are described in **Table S1**. *Mycoplasma capricolum* subsp. *capricolum (Mcap)* strain ATCC 27343 (California Kid^T^), *Mycoplasma mycoides* subsp. *capri* (*Mmc*) strain GM12, and *Mycoplasma mycoides* subsp. *mycoides* (*Mmm*) strain T1/44 were grown in SP5 medium^24^. *Mycoplasma capricolum* subsp. *capripneumoniae (Mccp)* strains 95043 and 14020 were grown at 37°C in modified Hayflick’s medium (m-Hayflick medium)^56^. *Mycoplasma gallisepticum* strain S6^T^ was grown at 37°C in m-Hayflick medium^56^. *Mycoplasma bovis* strain PG45^T^ and *Mycoplasma agalactiae* strain PG2^T^ were cultured in SP4 bovis medium^56^. All strains were incubated at 37°C under a 5% CO_2_ atmosphere. Media were supplemented with 10 µg.mL^-1^ of puromycin for *M. gallisepticum mnuA* mutant and with 100 µg.mL^-1^ of gentamycin for *M. bovis and M. agalactiae mnuA* mutants.

*Saccharomyces cerevisiae* strain VL6−48N (MATa, *his3*-Δ200, *trp1*-Δ1, *ura3*−52, *lys2*, *ade2*−101, *met14*) was grown at 30 °C in YPDA medium (Clontech Takara). Yeast transformants were selected in Synthetic Defined (SD) medium depleted for one or several amino-acids: SD-Trp, SD-Trp-Ura or SD-His-Leu (Clontech Takara).

*Escherichia coli* strain NEB5-α (NEB C2987) used for plasmid cloning were grown at 37 °C in lysogenic broth (LB) medium supplemented with 100 µg.mL^-1^ of ampicillin.

### Plasmids and oligonucleotides

The plasmids and oligonucleotides used in this study are described and reported in **Table S7.** Two plasmids were used for CRISPR-Cas9 editing. The p414-TEF 1p-Cas9-CYC1t plasmid (Cas9 expression) developed by DiCarlo *et al.*^57^, and several pgRNA plasmids derived from the p426-SNR52p-gRNA.AarI.Y-SUP4t described by Tsarmpopoulos *et al.*^58^ and based on the p426-SNR52p-gRNA.CAN1.Y-SUP4t plasmid also developed by DiCarlo *et al.*^57^. Plasmids used for PCR template amplification routinely used in the laboratory are named pMT85-pRS313-pSTetM-pSLacZ^24^, pVC604-pRS313-pSTetM-pSLacZ, and pMT85-pRS313-pGenta-pSLacZ (Personal communication from Dr. Carole Lartigue; Gourgues *et al.* unpublished). Specific plasmids for deaminase application were construct by Gibson assembly as described in Ipoutcha *et al.*, 2022^49^. More precisely pFRIT4.0-mnuA or pTi4.0_Sp_pmcda_NucGalli0070 was used for MnuA homologs inactivation in M. agalactiae or in M. gallisepticum respectively.

### Construction of gRNA plasmids for simple target deletion

gRNA targeting selected mycoplasma *loci* (**Table S7**) were designed using Benchling [Biology Software] (retrieved from https://benchling.com). Corresponding pgRNA plasmids were constructed following the protocol described in Tsarmpopoulos *et al.,*2016^58^. Briefly, the plasmid p426-SNR52p-gRNA.AarI.Y-SUP4t contains all the elements necessary for the expression of the gRNA in yeast^57, 58^. The spacer component of the gRNA can be swapped out by restriction of the plasmid using AarI, followed by ligation of annealed oligonucleotides pairs. The resulting plasmids are transformed in *E. coli* and sequence verified.

### Construction of gRNA plasmids (pgRNA) for double cutting and genome fragment capture

The cassettes allowing the expression of the gRNA_Mmm_*glpOKF* and the gRNA_RS1980 or gRNA_RS1995 were PCR amplified from single target gRNA plasmid. Amplicons were then cloned using Gibson Assembly Cloning Kit (NEB), producing four versions of pDT-gRNA-*glpOKF-*RS (A, B, C or D) plasmids. Those versions are different in consistence with the gRNA-RS (see **Table S7**).

### Base editing of mnuA to produce M. gallisepticum, M. bovis and M. agalactiae mutant strains

*M. bovis mnuA* mutant was previously produced and described in Ipoutcha *et al*. 2022^49^. *M. gallisepticum mnuA* mutant and *M. agalactiae mnuA* mutant were produced for this study following the same protocol. Difference was more precisely on the target design and plasmid construction (see **Table S7**). Rapidly, for each *Mycoplasma* species, transformants on the third passage were grown in selective media (dilution 1/100). When culture was in early logarithmic growth phase, fresh anhydrotetracyclin (aTC) in ETOH 50% was added until the stationary phase (final concentration at 0.5 µg.mL^-1^). Alternatively, direct induction during transformation could be performed. To do so, after 2 h incubation at 37°C, antibiotics (puromycin or gentamicin) were then added to the media for two hours. Then, base editor system was induced using fresh aTC (0.5 µg.mL^-1^) for 12 h to 15 h (overnight). Induced cultures were plated on selective media and incubated at 37°C with 5% CO2.

### *Mycoplasma* transformation protocol

*M. gallisepticum* and *M. agalactiae* were transformed with the pTi4.0_Sp_pmcda_NucGalli0070 or pFRIT4.0-mnuA plasmids using protocols described by Ipoutcha *et al.*, 2022^49, 59^ and Zhu *et al.*, 2020^60^ respectively. Transformants were selected on appropriate solid media containing antibiotics (10µg.ml^-1^ of puromycin for *M. gallisepticum* and with 100 µg.mL^-1^ of gentamycin for *M. agalactiae*). Isolated colonies were picked and cultured in liquid media supplemented with the same antibiotic selection during 3 passages.

### Assays for MnuA nuclease activity

The nuclease activity of wild-type *M. bovis* and *M. bovis mnuA* mutant was assessed and reported in Ipoutcha *et al.* 2022 following a protocol previously published^49^. Briefly, cells were grown to a late-log phase culture and after centrifugation at 7000 x *g* for 10 minutes at 10°C, were suspended in 500 µL of nuclease reaction buffer (25 mM Tris-HCl, pH 8.8, 10 mM CaCl_2_, 10 mM MgCl_2_). The cell preparations were co-incubated with plasmid DNA (used as control in CReasPy-Fusion experiments) or with linear double strand DNA (recombination template carrying yeast elements), for 5 or 60 minutes at 37°C. At each time point, 10 µL were sampled and the reaction were stopped by the addition of EDTA to a final concentration of 20 mM. Analysis was done by 1% TAE (Tris-Acetate 40 mM; EDTA 1 mM; pH 8.0) agarose gel electrophoresis migration of each sample mixed with 6X loading buffer (Gel Loading Dye Purple NEB). DNA degradation was appreciated after ethidium bromide coloration (2µg/mL final concentration) and UV exposure.

### Plasmid transformation in yeast

Yeast was transformed using the lithium acetate protocol optimized by Gietz *et al.*^61^. One μg of purified plasmid (pCas9 (TRP1) and/or pgRNA (URA3)) was used for each transformation, and transformants were selected for auxotrophy complementation in SD-Trp or SD-Trp-Ura.

### Construction of recombination templates

Recombination templates containing the yeast elements are produced by PCR amplification of the ARS/CEN/HIS/PSTetM, CEN/HIS/PSTetM, ARS/CEN/HIS/Genta and CEN/HIS/Genta *loci* from corresponding plasmid templates indicated in supplemental **Table S7**. Specific primers were designed for this purpose and PCR was done using the Advantage 2 Polymerase kit (Clontech). Complementary 60 bp-ends to each target sequence on all *Mycoplasmas* species genome used in this study, were added to the extremities of the cassettes by using 5′-tailed PCR primers.

### Yeast Fusion with *Mycoplasma* cells and recombination template

Yeast cells carrying the pCas9 and pgRNA plasmids were fused with mycoplasma cells (WT or mutants) as described by Karas et al^22, 28^. Briefly, 200 μL of yeast spheroplasts were mixed with 50µL of mycoplasma cells and 1 µg of recombination template containing the yeast elements. Bacterial cells were previously warmed up before the Fusion step at a fixing temperature of 49°C for all species. After transformation, the yeast cells were selected on SD-His-Leu solid agar plates containing 1 M of sorbitol, for 4 days at 30°C. Individual colonies were picked and streaked on SD-His-Leu plates and incubated 2 days at 30 °C. Then, one isolated colony per streak is patched on the same medium and incubated for 2 days at 30 °C.

### Screening of yeast transformants carrying mycoplasma genomes

Total genomic DNA was extracted from yeast transformants according to Kouprina and Larionov^25^. Positive clones were screened for both the presence of the *Mycoplasma* genome and the correct deletion of the target genes by PCR, using the Advantage 2 Polymerase kit (Clontech) and specific primers located on either side of the target locus. Yeast transformants were then screened for bacterial genome completeness by multiplex PCR using specific sets of PCR primers for each *Mycoplasma* species (**Table S7**). Each set is comprised of ten pairs of primers evenly distributed across the bacterial genomes allowing the simultaneous amplification of fragments ranging from ∼100 to ∼1000 bp, in ∼100 bp increments. Clones carrying mycoplasma genomes with no major rearrangements display a characteristic ten-bands or eleven-bands ladder profile with each primer set. The multiplex PCR are performed using the Qiagen Multiplex PCR Kit according to the manufacturer’s instructions. Yeast clones appearing positive by multiplex PCR are ultimately analyzed by restriction digestion and pulsed-field gel electrophoresis (PFGE) to assess the size of the mycoplasma chromosome. To do so, yeast cells were grown in SD-His media harvested, embedded in agarose plugs and lysed by treatments with zymolyase, proteinase K, and detergents to yield intact chromosomes using CHEF Mammalian Genomic Plug kit Bio-Rad. At this stage, yeast plugs carrying *Mycoplasma* genomes were treated slightly differently. Analysis by pulsed field electrophoresis consisted of a series of steps to remove the yeast DNA, and then to migrate the mycoplasma genome according to its size (linearization or digestion of the genome in several pieces). For most of the mycoplasma genomes cloned in yeast and analyzed in this work, the agarose plugs were treated with a cocktail of restriction enzymes (AsiSI, FseI, RsrII) hydrolyzing exclusively yeast genomic DNA. Then, there were loaded onto a conventional electrophoresis gel (1% (w/v) agarose, TAE1 X, 120V, 120min.) in order to eliminate a large part of the yeast DNA and to avoid that the latter masks the digestion profile of the bacterial genome that we wish to analyze. An exception to this procedure was the analysis of half-plugs containing DNA from yeast transformed with a *M. gallisepticum* genome. This elimination step is performed directly with a first PFGE migration (1% agarose, 0.5×TBE) during 24h, with a switch time of 50−90 s, at 6 V cm−1, an angle of 120° and a temperature of 14°C. Then, after the electro-removal of the yeast linear chromosomes, the DNA remaining in plugs was restricted with BssHII (for *M. capricolum* subsp. *capricolum*, *M. capricolum* subsp. *capripneumoniae*, *M. mycoides* subsp. *mycoides*), SfoI (for *M. mycoides* subsp. *capri*), EagI (for *M. mycoides* subsp. *mycoides*), SacII (for *M. gallisepticum*) and AscI (for *M. agalactiae*), and submitted to PFGE. Pulse times are ramped from 60s to 120 s for 24 h at 6 V cm−1. Agarose gels were stained with SYBR Gold Nucleic Acid Gel Stain to (Invitrogen) and PFGE patterns were scanned using the Vilbert Lourmat E-BOX VX2 Complete Imaging system.

### Transplantation of mycoplasma genome cloned in yeast into a recipient cell

Following the protocol described in Lartigue *et al.* 2007 and 2009 and improved in Labroussaa *et al.* 2016.

### Whole genome sequencing of mycoplasma transplants

Genomic DNA of *Mcap* CK^T^ cl12.1, *Mmc* GM12 cl 5.1 and *Mccp* Tanzanie cl4.2 transplants was extracted from a 10 mL culture using the Qiagen Genomic-Tips 100/G kit. Genome sequencing was performed by the Genome Transcriptome Facility of Bordeaux. Long reads were produced using a GridION device (Oxford Nanopore) and short reads a MiSeq device (Illumina). For the *Mcap* cl12.1 transplant, ONT sequencing generated 31,638 reads (mean read length: 21,911 bp) and Illumina 366,632 read pairs. For the *Mmc* cl5.1 transplant we obtained 21,267 ONT long-reads (mean read length: 23,569 bp) and 190,934 Illumina short-read pairs. For the newly *Mccp* WT Tanzanian referee genome (sequence for this work and publish in association) and the *Mccp* cl4.2 transplant, ONT sequencing generated respectively 11,051 and 20,650 reads (mean read length: 26,332 and 18,015 bp), and we obtained 294,650 and 240,318 Illumina short-read pairs. All the analysis were performed using Galaxy (https://usegalaxy.eu/). Genome assembly was performed using the following steps: Nanopore reads were filtered using Filter FASTQ (V 1.1.5, Minimum size 45000 bp), assembled using Flye Assembly (V 2.6), polished using 4 rounds of Pilon (1.20.1) combined with Illumina short reads. These last are combined and sorted out with Trimmomatic (V 0.38.1; Sliding Window 10, 20; Drop read below minimal length of 250), and the quality was checked with Fastqc (Version 0.11.8). Assembled genome was compared to each species corresponding reference genome: *Mcap* CK^T^ (CP000123.1), *Mmc* GM12 (CP001668.1) and, *Mccp* Tanzania (specifically sequence for this work and published in association (CP121686.1)). In a second part, mutations were detected after mapping the Illumina reads onto the referees genomes cited above using a second pipeline : Illumina reads were trimmed using Trimmomatic (V 0.38.1; Sliding Window 10, 20; Drop read below minimal length of 250), mapped using BWA-MEM (V 0.7.17.1), Samtools sort (V 2.0.3), and MPileup (V 2.1.1), and variants detected using VarScan mpileup (V 2.4.3.1; Minimum coverage 30, Minimum supporting read 20, Minimum Base quality 30, Minimum variant allele frequency 0.8, Minimum homozygous variants 0.75). Results of these analyses are shown in **Table S11**.

## ASSOCIATED CONTENT

### Supporting Information

Supplementary Material: this pdf file includes Supplementary Figures S1 to S6. Supplementary Table S1 to S11 (excel files for Table S1, S5, S7 and S11).

### Data availability

We declare that the data supporting the findings of this study are available within the paper and its supplemental material. The sequences from this study are available from the NCBI under SRA accession no. PRJNA910329. The updated version of the *Mccp* strain 14020 Tanzania genome was deposited in GenBank under the BioProject PRJNA940743, the BioSample SAMN33578400 and the accession number CP121686.1.

### Author Contributions

Conceptualization, G.Gu, PS-P, C.L.

Formal Analysis, G.Gu, C.L.

Funding acquisition, M.J, PS-P, C.L. Investigation, G.Gu, C.L.

Methodology, G.Gu, G.Go, F.R, T.I and C.L.

Whole genome sequence analysis, G.Gu, T.I, PS-P and C.L.

Resources, L.M-S.

Supervision, A.B, PS-P, C.L.

Validation, G.Gu, G.Go, F.R, T.I and C.L.

Visualization, G.Gu, G.Go, F.R, T.I and C.L.

Writing – original draft, G.Gu, A.B, C.L.

Writing – review & editing: all authors approved the final version of the manuscript.

### Notes

The authors declare no competing financial interests.

## ACKNOWLEDGMENT

The authors thank Christophe Boury and Erwan Guichoux members of the Genome Transcriptome Facility of Bordeaux (https://pgtb.fr/, Grants from Investissements d’avenir, Convention attributive d’aide EquipEx Xyloforest, ANR-10-EQPX-16-01), for their help in genome sequencing performance. We also thank the UMR CIRAD-INRAE ASTRE from Montpellier for providing the *Mccp* Niger and Tanzania strains; and the UMR IHAP INRAE-ENVT from Toulouse, for providing the *M. bovis* PG45^T^ strain and the *M. agalactiae* PG2^T^.

