## Supplemental Information for "Combining fusion of cells with CRISPR-Cas9 editing for the cloning of large DNA fragments or complete bacterial genomes in yeast"


**Authors and affiliations**

Gabrielle Guesdon[1], Géraldine Gourgues[1], Fabien Rideau[1], Thomas Ipoutcha[1], Lucía Manso-Silván[2,3], Matthieu Jules[4], Pascal Sirand-Pugnet[1], Alain Blanchard[1] and Carole Lartigue[1§]

[1] Univ. Bordeaux, INRAE, Biologie du Fruit et Pathologie, UMR 1332, F-33140 Villenave d'Ornon, France

[2] CIRAD, UMR ASTRE, F-34398 Montpellier, France.

[3] ASTRE, Univ Montpellier, CIRAD, INRAE, F-34398, Montpellier, France.

[4] Université Paris-Saclay, INRAE, AgroParisTech, Micalis Institute, 78350, Jouy-en-Josas, France.

[§] **Corresponding author:**

INRAE, Equipe Mollicutes, UMR 1332 BFP, Bâtiment IBVM - A4, 71, Avenue Edouard Bourlaux, CS 20032, F-33882 VILLENAVE D'ORNON CEDEX, France


**This file includes**

Supplementary Figures S1 to S6

Supplementary Tables S1 to S11

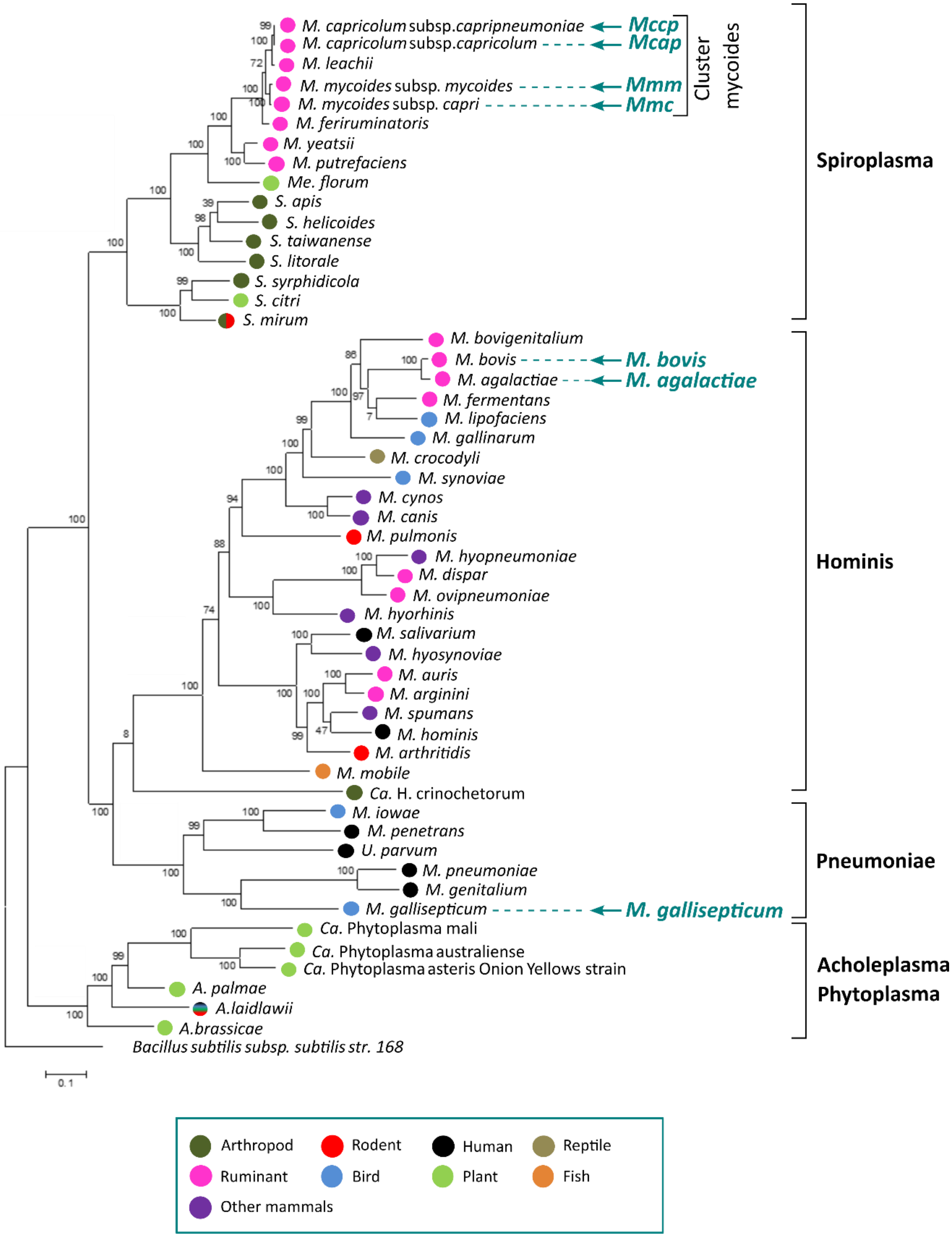


**Figure S1: Phylogenetic tree of Mollicutes.** The phylogenetic tree was generated using concatenated multiple sequence alignments of selected 62 orthologous protein involved in translation (Maximum Likelihood method). The major phylogenetic groups Spiroplasma, Hominis, Pneumoniae, Acholeplasma/Phytoplasma are indicated, as well as the mycoides cluster. The seven species used for the CReasPy-Fusion cloning method are indicated by turquoise arrows. The hosts of each species are indicated by colored circles following the color code specified in the box. The species *Bacillus subtilis* was used as an outgroup to root the phylogenetic tree.

Guesdon *et al*,

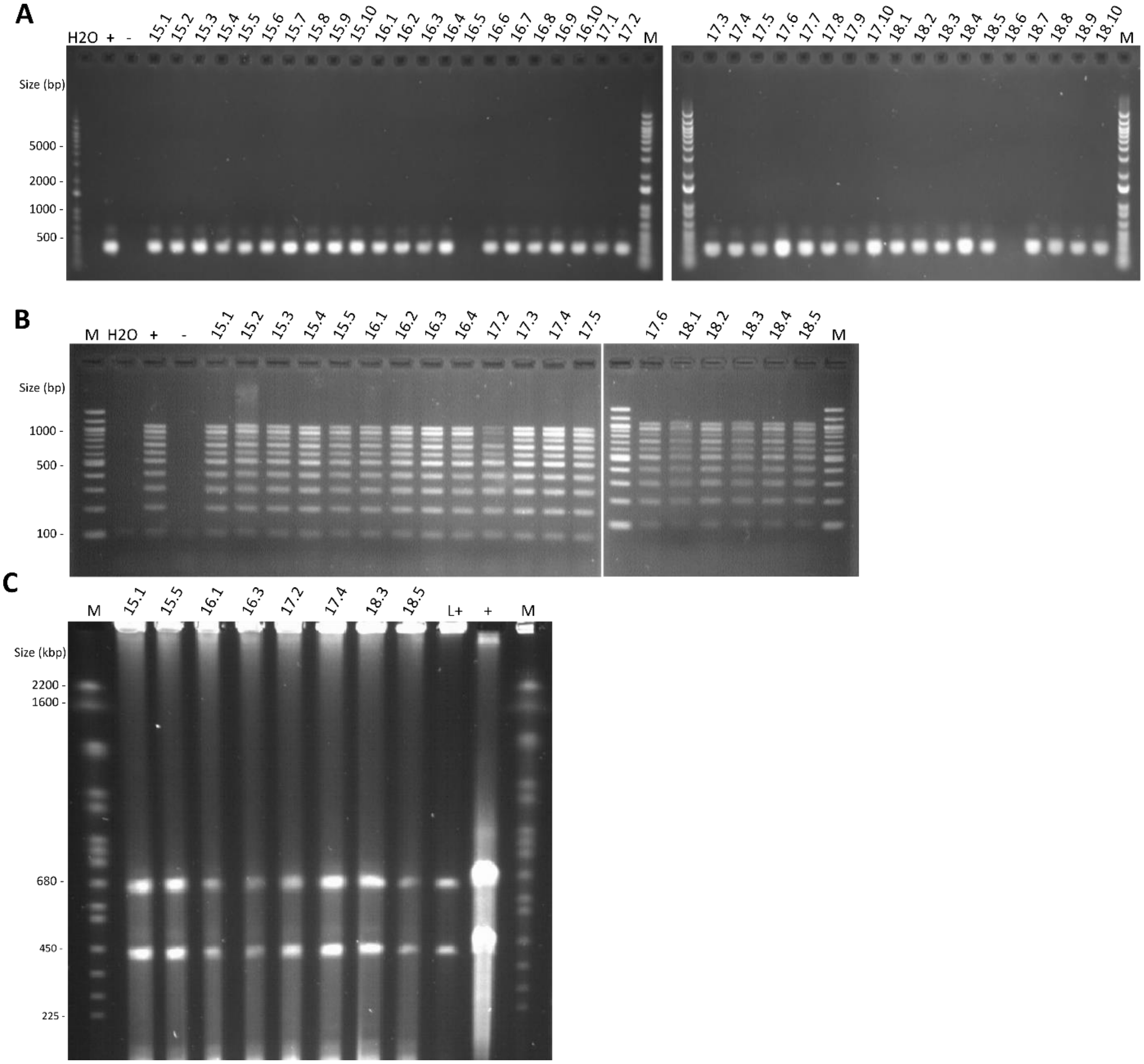


**Figure S2: Screening of yeast transformants carrying the entire *McapΔRE* genomes after fusion.**
Results shown here represent the analysis performed for evaluating the fusion method for cloning genome from *McapΔRE*. Results of the screening by PCR simplex, PCR multiplex and PFGE performed are shown in the panels A, B and C, respectively. “M”: Simplex PCR = DNA Ladder 1kb+ Invitrogen (100-12,000 bp); Multiplex PCR: DNA Ladder 100bp NEB (100-1,517 bp) ; PFGE = *S. cerevisiae* chromosomal DNA ladder Bio-Rad (225-2,200 kbp). “H2O”: negative control without DNA. “+”: positive control with *McapΔRE* gDNA. “L+”: gDNA positive control of yeast transformant carrying *McapΔRE* genome. “-“: negative control with *S. cerevisiae* VL6-48N gDNA. **(A)** The presence of *McapΔRE* genome is verified by PCR analysis using specific primers (MCAPCK_F1/R1) generating a 272 bp amplicon as obtained with the positive control. **(B)** The completeness of the *McapΔRE* genome cloned in yeast was assessed by multiplex PCR using a set of primers distributed around the genome (from 100 bp to 1,000 bp). Clones carrying genomes without major rearrangement displayed a ten-band profile identical to the one obtained in the positive control. **(C)** The size of the genome cloned in yeast was assessed by enzymatic restriction and Pulsed Field Gel Electrophoresis (PFGE). Digestion of the bacterial genome with the restriction enzyme BssHI is expected to generate two DNA fragments of 626 and 383 kbp.

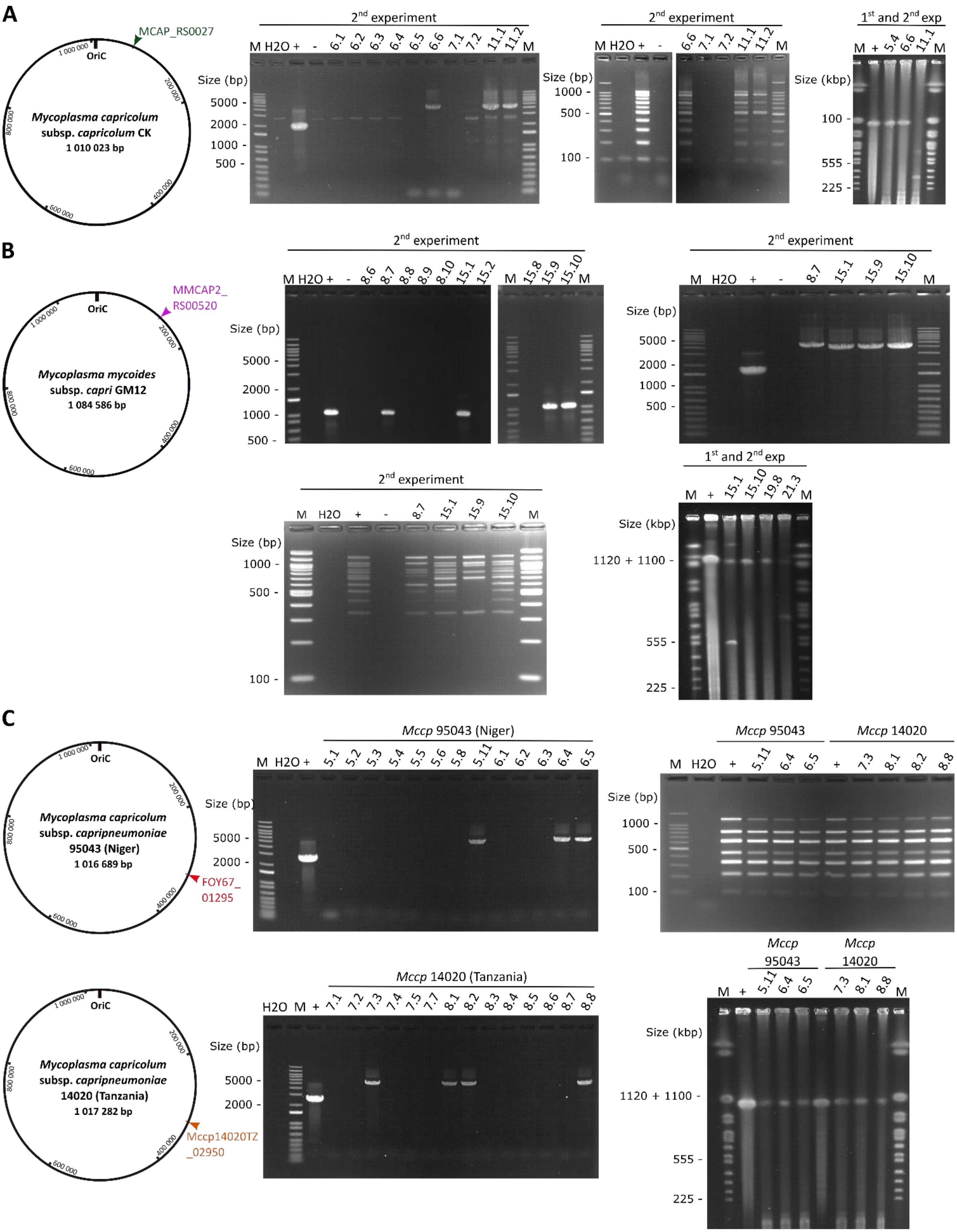


Guesdon *et al*,

**Figure S3: Screening of yeast transformants carrying entire *Mcap*, *Mmc* or *Mccp* genomes after CReasPy-Fusion.** Results shown here represent the analysis performed for evaluating CReasPy-Fusion method for cloning the genome from *Mycoplasma capricolum* subsp *capricolum* CK (*Mcap* CK), *Mycoplasma mycoides* subsp *capri* GM12 (*Mmc* GM12) and *Mycoplasma capricolum* subsp *capripneumoniae* strain 95043 Niger and strain 14020 Tanzania (*Mccp* 95043 Niger or *Mccp* Tanzania). For each *Mycoplasma* species, the genome map with location of targeted loci is indicated by coloured arrow heads and the screening by PCR simplex, PCR multiplex and PFGE performed is summarized. "M": Simplex PCR = DNA Ladder 1kb+ Invitrogen (100-12000 bp); Multiplex PCR: DNA Ladder 100 bp NEB (100-1517 bp); PFGE = *S. cerevisiae* chromosomal DNA ladder Bio-Rad (225-2200 kpb). "H2O": negative control without DNA. "+": positive control with *Mcap*, *Mmc* or *Mccp* gDNA. "-": negative control with *S. cerevisiae* VL6-48N gDNA. **(A)** Left to the right: the MCAP_RS00270 deletion is validated by PCR analysis using specific primers flanking the target gene. Positive transformants were validated with a 4626 pb or 4905 pb amplicon (+/- ARS) instead of the 1,966 pb WT expected loci amplicon. The completeness of the *Mcap* CK genome cloned in yeast was assessed by multiplex PCR using a set of primers distributed around the genome (from 100 bp to 1,000 bp). Clones carrying genomes without major rearrangement displayed a ten-band profile identical to the one obtained in the positive control. The size of the *Mcap* CK genome cloned in yeast was assessed by enzymatic restriction and Pulsed Field Gel Electrophoresis (PFGE). Digestion of the bacterial genome with the restriction enzyme BssHII should generate one linear DNA fragment of 1,010 kbp. **(B)** Left to the right in first line: the presence of *Mmc* GM12 genome is checked by PCR analysis using specific primers generated a 1,010 bp amplicon as obtained in the positive control and the MMCAP2_RS00520 deletion is validated by PCR analysis using specific primers flanking the target gene. Positive transformants were validated with a 4,280 pb or 4,757 pb amplicon (+/- ARS) instead of the 1,719 pb WT expected loci amplicon. Left to the right in second line: the completeness of the *Mmc* GM12 genome cloned in yeast was assessed by multiplex PCR using a set of primers distributed around the genome (from 377 bp to 1,149 bp). Clones carrying genomes without major rearrangement displayed an eleven-band profile identical to the one obtained in the positive control. The size of the *Mmc* GM12 genome cloned in yeast was assessed by enzymatic restriction and Pulsed Field Gel Electrophoresis (PFGE). Digestion of the bacterial genome with the restriction enzyme SfoI should generate one linear DNA fragment of 1,084kbp. **(C)** Respectively for *Mccp* 95043 (Niger) and *Mccp* 14020 (Tanzania), the FOY67_01295 and the Mccp14020TZ_02950 deletion are validated by PCR analysis using specific primers flanking the target gene(s). Positive transformants were validated with a 5,044 pb amplicon instead of the 2,501 pb WT expected loci amplicon. The completeness of the *Mccp* 95043 or 14020 genome cloned in yeast was assessed by multiplex PCR using a set of primers distributed around the genome (from 216 bp to 1,205 bp). Clones carrying genomes without major rearrangement displayed a seven-bands profile identical to the one obtained in the positive control. For both strains. The size of the *Mccp* genome cloned in yeast was assessed by enzymatic restriction and Pulsed Field Gel Electrophoresis (PFGE). Digestion of the bacterial genome with the restriction enzyme BssHII should generate one linear DNA fragment of 1,016 kbp.

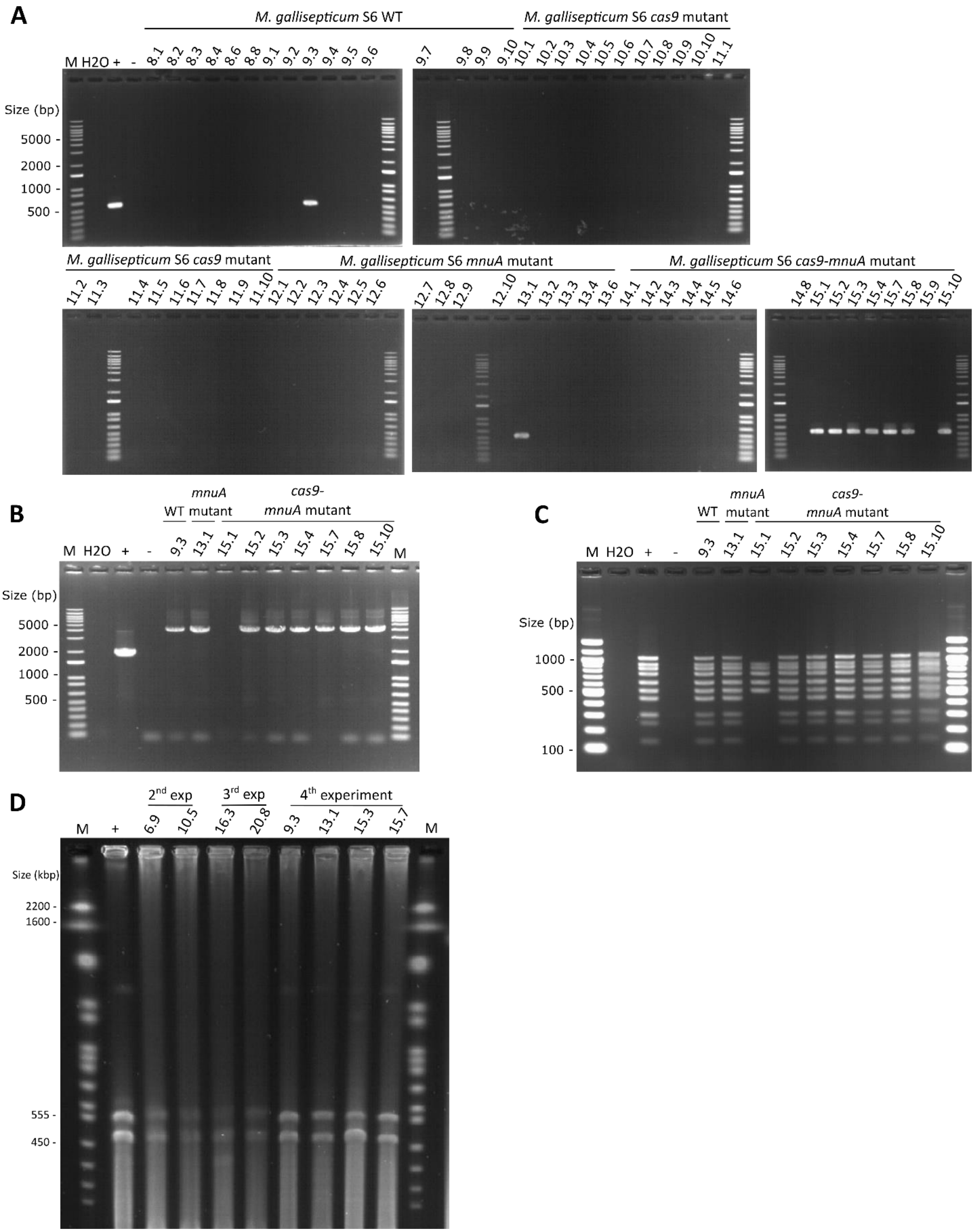


*Guesdon et al,*

**Figure S4: Screening of yeast transformants carrying the entire *M. gallisepticum* genome after CReasPy-Fusion.** In this figure (panels A, B and C), an example of the screening performed for the 4$^{th}$ experiment is shown. **(A)** The presence of *M. gallisepticum* S6 genome (WT or mutants) is checked by PCR analysis using specific primers generating a 528 bp amplicon as obtained in the positive control. **(B)** The GCW_RS01695 deletion is validated by PCR analysis using specific primers flanking the target gene. Positive transformants were validated with a 4,503 pb amplicon instead of the 1,952 pb for the WT expected loci amplicon. **(C)** The completeness of the *M. gallisepticum* S6 (WT or mutants) genome cloned in yeast was assessed by multiplex PCR using a set of primers distributed around the genome (from 137 bp to 1,038 bp). Clones carrying genomes without major rearrangement displayed a ten bands profile identical to the one obtained in the positive control. **(D)** The size of the entire *M. gallisepticum* S6 (WT or mutants) genome cloned in yeast was assessed by enzymatic restriction and PFGE. Digestion of the bacterial genome with the restriction enzyme SacII should generate two DNA fragments of 536 and 449 kbp. "M": Simplex PCR = DNA Ladder 1kb+ Invitrogen (100-12,000 bp); Multiplex PCR: DNA Ladder 100bp NEB (100-1,517 bp); PFGE = *S. cerevisiae* chromosomal DNA ladder Bio-Rad (225-2,200 kbp). "H2O": negative control without DNA.

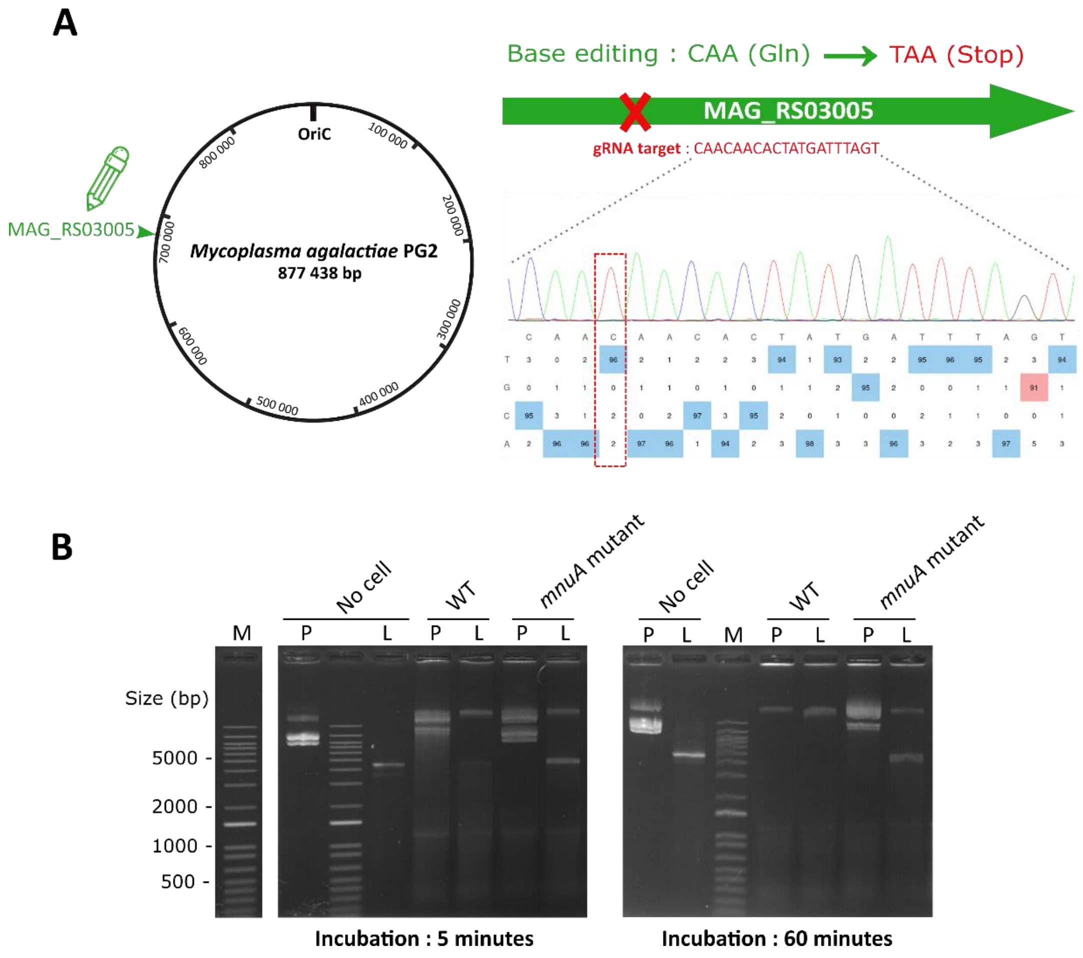


**Figure S5: Inactivation of the nuclease membrane MnuA homolog in *M. agalactiae* PG2 genome.**

**(A)** Map of *M.agalactiae* PG2 genome. The location of the targeted locus (MAG_RS03005) is indicated by colored arrow head. Base editing modification of the same locus gene (membrane nuclease homolog) is indicated by a green pencil scheme: the gRNA was design to edit CAA glycine codon into a TAA stop codon involving inactivation of the target gene. Chromatogram result associated with the EditR software analysis software analysis (https://moriaritylab.shinyapps.io/editr_v10/) are represented and the "T peak" instead of a "C peak" change is indicated with red rectangle dotted line **(B)** Nuclease activity assay is shown for both wild-type and *M. agalactiae* mutants after 5 minutes (left) or 60 minutes (right) of incubation. "M": DNA Ladder 1kb+ Invitrogen (100-12,000 bp); "P": plasmid DNA ; "L" : linear DNA.

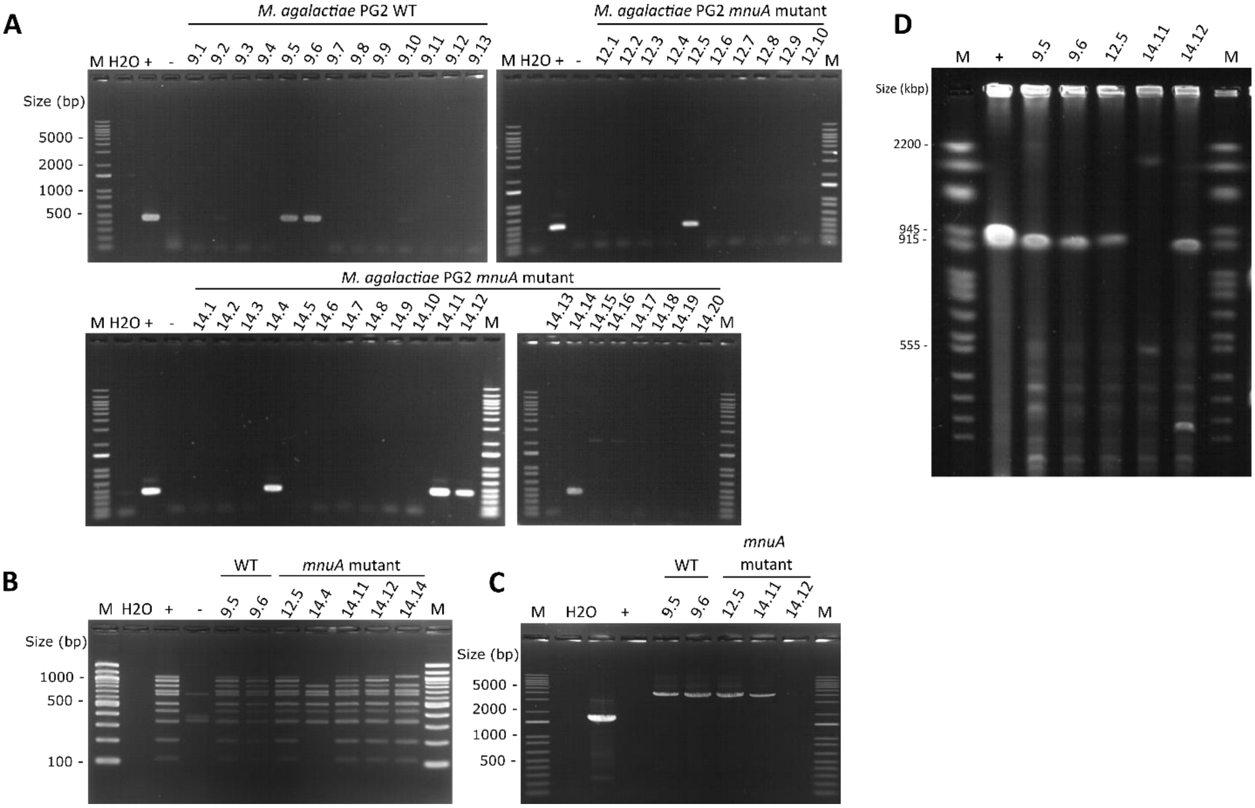


**Figure S6: Screening of yeast transformants carrying the entire *M. agalactiae* genome after CReasPy-Fusion. (A)** The presence of *M. agalactiae* PG2 genome (WT or mutants) is checked by PCR analysis using specific primers generating a 425 bp amplicon as obtained in the positive control. **(B)** The completeness of the *M. agalactiae* PG2 (WT or mutants) genome cloned in yeast was assessed by multiplex PCR using a set of primers distributed around the genome (from 111 bp to 1,037 bp). Clones carrying genomes without major rearrangement displayed a ten-band profile identical to the one obtained in the positive control. **(C)** The MAG_RS03005 deletion is validated by PCR analysis using specific primers flanking the target gene. Positive transformants were validated with a 4364 or 4841 pb amplicon (+/-ARS) instead of the 1,712 pb expected amplicon for the WT genome. **(D)** The size of the entire *M. agalactiae* PG2 (WT or mutants) genome cloned in yeast was assessed by enzymatic restriction and Pulsed Field Gel Electrophoresis (PFGE). Digestion of the bacterial genome with the restriction enzyme AscI should generate a single linear DNA fragment of 877 kbp. "M": Simplex PCR = DNA Ladder 1kb+ Invitrogen (100-12,000 bp); Multiplex PCR: DNA Ladder 100bp NEB (100-1,517 bp); PFGE = *S. cerevisiae* chromosomal DNA ladder Bio-Rad (225-2,200 kbp). "H2O": negative control without DNA. "+": positive control with *M. agalactiae* gDNA. "-": negative control with *S. cerevisiae* VL6-48N gDNA.

**Table S1: Summary of mycoplasma species and strains used for CReasPy-Fusion experiments** **(**Excel document).

**Table S2: In-yeast cloning of the *McapΔRE* genome by cell fusion**

| Yeast spheroplast | *M. capricolum* subsp. *capricolum* ΔRE | UFC | Positive clones / Analyzed clones[a] | | |
|---|---|---|---|---|---|
| | | | Simplex PCR | Multiplex PCR[b] | PFGE[b] |
| 100µL | 50µL | 88 | 10/10 | 5/5 | 2/2 |
| | 100µL | 97 | 9/10 | 4/4 | 2/2 |
| 200µL | 50µL | 202 | 10/10 | 5/5 | 2/2 |
| | 100µL | 185 | 9/10 | 5/5 | 2/2 |

[a] One Fusion experiment was performed. The number of yeast transformants analyzed by simplex PCR, multiplex PCR and PFGE is reported, as well as the number of positive clones obtained.
[b]Multiplex PCR and PFGE analysis were not performed for all the positive clones but only for a sample.

**Table S3: Genome transplantation (GT) experiment conducted with mycoplasma genomes *Mcap* CK, *Mmc* GM12 and *Mccp* (Niger and Tanzania)**

| | Donor genome | CFU | Positive clones / Analyzed clones[a] | |
|---|---|---|---|---|
| | | | Multiplex PCR | Sequencing |
| **A- *Mcap* GT** | VL6-48N x *Mcap*CKΔ0050(RE-CCATC) cl5.4 | 3 | 3/3 | cl12.1 selected |
| | VL6-48N x *Mcap*CKΔ0050(RE-CCATC) cl6.6 | 0 | - | - |
| **B- *Mmc* GT** | VL6-48N x *Mmc*GM12Δ0100(RE-CCATC) cl15.1 | 3 | 2/2 | cl5.1 selected |
| | VL6-48N x *Mmc*GM12Δ0100(RE-CCATC) cl15.10 | 0 | - | - |
| | VL6-48N x *Mmc*GM12Δ0100(RE-CCATC) cl19.8 | 9 | 4/4[b] | - |
| | VL6-48N x *Mmc*GM12Δ0100(RE-CCATC) cl21.3 | 0 | - | - |
| **C- *Mccp* GT** | VL6-48N x *Mccp*95043NigerΔ01295(pepS41) cl5.11 | 28 | 4/4[b] | - |
| | VL6-48N x *Mccp*95043NigerΔ01295(pepS41) cl6.4 | 25 | 4/4[b] | - |
| | VL6-48N x *Mccp*95043NigerΔ01295(pepS41) cl6.5 | 33 | 4/4[b] | - |
| | VL6-48N x *Mccp*1420TanzaniaΔ02950(pepS41) cl7.3 | 2 | 1/1 | cl4.2 selected |
| | VL6-48N x *Mccp*1420TanzaniaΔ02950(pepS41) cl8.1 | 3 | 2/2 | - |
| | VL6-48N x *Mccp*1420TanzaniaΔ02950(pepS41) cl8.2 | - | - | - |
| | VL6-48N x *Mccp*1420TanzaniaΔ02950(pepS41) cl8.8 | 1 | 1/1 | - |

[a] The number of yeast transformants analyzed by simplex PCR, multiplex PCR and PFGE is reported, as well as the number of positive clones obtained.
[b]Multiplex PCR analysis were not performed for all the positive clones but only for a sample.

**Table S4: *M. gallisepticum* WT genome cloning first assay by CReasPy-Fusion**

| Condition | Yeast | Spheroplast volume | Plasmid[§] or recombination template[ǂ] | DNA quantity | *M. gallisepticum* cell volume | CFU | Positive clones / Analyzed clones[a] | |
|---|---|---|---|---|---|---|---|---|
| | | | | | | | Analyzed | Simplex PCR |
| **1** | VL6-48N-pCas9-pARNg1 | 200 µL | - | - | - | 0 | - | - |
| **2** | | | pMT85[§] | 300 ng | - | >1000[b] | - | - |
| **3** | | | | | 100 µL | 1[b] | - | - |
| **4** | | | ARS-CEN-HIS-CysP[ǂ] | 1 µg | - | >300 | - | - |
| **5** | | | | | 25 µL | 80 | 20 | 0/20 |
| **6** | | | | | 50 µL | 10 | 10 | 0/10 |
| **7** | | | | | 100 µL | 0 | - | - |
| **8** | | | CEN-HIS-CysP[ǂ] | | - | 34 | - | - |
| **9** | | | | | 25 µL | 7 | 7 | 0/7 |
| **10** | | | | | 50 µL | 1 | 1 | 0/1 |
| **11** | | | | | 100 µL | 0 | - | - |
| **12** | VL6-48N-pCas9-pARNg2 | 200 µL | - | - | - | 0 | - | - |
| **13** | | | pMT85[§] | 300 ng | - | >1000 | - | - |
| **14** | | | | | 100 µL | 20 | - | - |
| **15** | | | ARS-CEN-HIS-CysP[ǂ] | 1 µg | - | >300 | - | - |
| **16** | | | | | 25 µL | 93 | 20 | 0/20 |
| **17** | | | | | 50 µL | 26 | 20 | 0/20 |
| **18** | | | | | 100 µL | 0 | - | - |
| **19** | | | CEN-HIS-CysP[ǂz] | | - | 45 | - | - |
| **20** | | | | | 25 µL | 29 | 20 | 0/20 |
| **21** | | | | | 50 µL | 10 | 10 | 0/10 |
| **22** | | | | | 100 µL | 0 | - | - |
| **23** | - | - | - | - | 100µL | 0 | - | - |

[a] The number of yeast transformants analyzed by simplex PCR, multiplex PCR and PFGE is reported, as well as the number of positive clones obtained.
[b]For control conditions (2 and 3) corresponding of plasmid (pMT85) transformation control protocol, the number of yeast transformants drastically decreased in addition of *M. gallisepticum* cells (>1000 vs 1 CFU).

**Table S5: Summary of restriction modification systems documented for mycoplasma species and strains used in CReasPy-Fusion experiments**
(Excel document).

**Table S6: *M. gallisepticum mnuA-cas9* mutant genome cloning assay by CReasPy-Fusion**

| | Experimental condition | CFU | Positive clones / Analyzed clones[a] | |
|---|---|---|---|---|
| | | | Simplex PCR | Multiplex PCR |
| **Controls** | Plasmid | >1000[b] | - | - |
| | Plasmid + *M. gallisepticum* | >1000[b] | - | - |
| | Plasmid + *M. gallisepticum* 49°C | >1000[b] | - | - |
| **1** | Resuspension Buffer | 86 | 2/10 | 2/2 |
| **2** | Resuspension Buffer + EDTA | 97 | 0/9 | - |
| **3** | Resuspension Buffer + denatured ssDNA | 1 | - | - |
| **4** | Resuspension Buffer + needle and filter treatment | 101 | 0/10 | - |
| **5** | Resuspension Buffer – 49°C | 106 | 2/10 | 2/2 |
| **6** | Resuspension Buffer + EDTA – 49°C | 92 | 2/9 | 2/2 |
| **7** | Resuspension Buffer + denatured ssDNA – 49°C | 4 | 0/3 | - |
| **8** | Resuspension Buffer + needle and filter treatment – 49°C | 120 | 1/10 | 1/1 |
| **9** | HBSS | 65 | 3/10 | 3/3 |
| **10** | HBSS + EDTA | 62 | 0/7 | - |
| **11** | HBSS + denatured ssDNA | 4 | 0/4 | - |
| **12** | HBSS + needle and filter treatment | 114 | 0/9 | - |
| **13** | HBSS – 49°C | 45 | 1/6 | 1/1 |
| **14** | HBSS + EDTA – 49°C | 48 | 0/7 | - |
| **15** | HBSS + denatured ssDNA – 49°C | 4 | 0/3 | - |
| **16** | HBSS + needle and filter treatment – 49°C | 87 | 0/10 | - |
| **Total** | - | 1036 | 11/117 | 11/117 |

[a]The number of yeast transformants analyzed by simplex PCR and multiplex PCR is reported, as well as the number of positive clones obtained.
[b]For control conditions corresponding of plasmid (pMT85) transformation control protocol, the number of yeast transformants remained identical in addition or not, of *M. gallisepticum mnuA* mutant cells (>1000).

**Table S7**: **Primers and plasmids used in this study**
(Excel document).

**Table S8:** ***M. bovis*** **WT genome cloning first assay by CReasPy-Fusion**

| Condition | Yeast | Spheroplast volume | Plasmid§ or recombination templateǂ | DNA quantity | *M. bovis* cell volume | CFU | Positive clones / Analyzed clones[a] | |
|---|---|---|---|---|---|---|---|---|
| | | | | | | | Analyzed | Simplex PCR |
| **1** | VL6-48N-pCas9-pARNg1 | 200 µL | - | - | - | 0 | - | - |
| **2** | | | pMT85§ | 300 ng | - | >500 | - | - |
| **3** | | | | | 100 µL | 164 | - | - |
| **4** | | | ARS-CEN-HIS-0215ǂ | 1 µg | - | >300 | - | - |
| **5** | | | | | 25 µL | 145 | 10 | 0/10 |
| **6** | | | | | 50 µL | 37 | 10 | 0/10 |
| **7** | | | | | 100 µL | 1 | 1 | 0/1 |
| **8** | | | CEN-HIS-0215ǂ | | - | >300 | - | - |
| **9** | | | | | 25 µL | 170 | 10 | 0/10 |
| **10** | | | | | 50 µL | 25 | 10 | 0/10 |
| **11** | | | | | 100 µL | 1 | 1 | 0/1 |
| **12** | VL6-48N-pCas9-pARNg2 | 200 µL | - | - | - | 0 | - | - |
| **13** | | | pMT85§ | 300 ng | - | >1000 | - | - |
| **14** | | | | | 100 µL | 137 | - | - |
| **15** | | | ARS-CEN-HIS-0215ǂ | 1 µg | - | >300 | - | - |
| **16** | | | | | 25 µL | 90 | 10 | 0/10 |
| **17** | | | | | 50 µL | 14 | 10 | 0/10 |
| **18** | | | | | 100 µL | 0 | - | - |
| **19** | | | CEN-HIS-0215ǂ | | - | >300 | - | - |
| **20** | | | | | 25 µL | 139 | 10 | 0/10 |
| **21** | | | | | 50 µL | 60 | 10 | 0/10 |
| **22** | | | | | 100 µL | 58 | 10 | 0/10 |
| **23** | - | - | - | - | 100µL | 0 | - | - |

[a] The number of yeast transformants analyzed by simplex PCR and multiplex PCR is reported, as well as the number of positive clones obtained.

**Table S9: *M. bovis mnuA* mutant genome cloning second assay by CReasPy-Fusion**

| Condition | Yeast | Plasmid[§] or recombination template[ǂ] | DNA quantity | *M. bovis* cell | Experimental condition | CFU | Positive clones / Analyzed clones[a] | |
|---|---|---|---|---|---|---|---|---|
| | | | | | | | Analyzed | Simplex PCR |
| **1** | VL6-48N-pCas9-pARNg1 | - | - | - | - | 0 | - | - |
| **2** | | pMT85[§] | 300 ng | - | - | >1000 | - | - |
| **3** | | | | 50 µL | Resuspension Buffer | >300 | - | - |
| **4** | | CEN-HIS-0215[ǂ] | 1 µg | - | - | >300 | - | - |
| **5** | | | | 50 µL | Resuspension Buffer | 153 | 20 | 0/20 |
| **6** | | | | | Resuspension Buffer + EDTA | 94 | 20 | 0/20 |
| **7** | | | | | dPBS | 23 | 20 | 0/20 |
| **8** | | | | | dPBS + EDTA | 24 | 20 | 0/20 |
| **9** | | | | | Resuspension Buffer – 49°C | 155 | 20 | 0/20 |
| **10** | | | | | Resuspension Buffer + EDTA – 49°C | 90 | 20 | 0/20 |
| **11** | | | | | dPBS – 49°C | 76 | 20 | 0/20 |
| **12** | | | | | dPBS + EDTA – 49°C | 196 | 20 | 0/20 |
| **13** | VL6-48N-pCas9-pARNg2 | - | - | - | - | 0 | - | - |
| **14** | | pMT85[§] | 300 ng | - | - | >1000 | - | - |
| **15** | | | | 50 µL | - | >300 | - | - |
| **16** | | CEN-HIS-0215[ǂ] | 1 µg | - | - | >300 | - | - |
| **17** | | | | 50 µL | Resuspension Buffer | 121 | 20 | 0/20 |
| **18** | | | | | Resuspension Buffer + EDTA | 48 | 20 | 0/20 |
| **19** | | | | | dPBS | 28 | 20 | 0/20 |
| **20** | | | | | dPBS + EDTA | 21 | 20 | 0/20 |
| **21** | | | | | Resuspension Buffer – 49°C | 55 | 20 | 0/20 |
| **22** | | | | | Resuspension Buffer + EDTA – 49°C | 22 | 20 | 0/20 |
| **23** | | | | | dPBS – 49°C | 47 | 20 | 0/20 |
| **24** | | | | | dPBS + EDTA – 49°C | 261 | 20 | 0/20 |
| **25** | - | - | - | 50µL | Resuspension Buffer | 0 | - | - |

[a] The number of yeast transformants analyzed by simplex PCR and multiplex PCR is reported, as well as the number of positive clones obtained.

**Table S10: *M. agalactiae* WT versus *mnuA* mutant genome cloning assay by CReasPy-Fusion**

| Condition | Yeast | Plasmid[§] or recombination template[ǂ] | DNA quantity | *M. agalactiae* cell | Experimental condition | CFU | Positive clones / Analyzed clones[a] | | | |
|---|---|---|---|---|---|---|---|---|---|---|
| | | | | | | | Analyzed | Simplex PCR | Multiplex PCR | PFGE |
| **1** | VL6-48N-pCas9-pARNg1+pARNg2 | - | - | - | - | 0 | - | - | - | - |
| **2** | | pMT85[§] | 300 ng | - | - | >1000[b] | - | - | - | - |
| **3** | | | | WT | Resuspension Buffer | 1[b] | - | - | - | - |
| **4** | | | | *mnuA* mutant | | >1000[b] | - | - | - | - |
| **5** | | | | WT | Resuspension Buffer – 49°C | 20 | - | - | - | - |
| **6** | | | | *mnuA* mutant | | >1000 | - | - | - | - |
| **7** | | CEN-HIS-0215[ǂ] | 1 µg | - | - | >500 | - | - | - | - |
| **8** | | | | WT | Resuspension Buffer | 3 | 3 | 0/3 | - | - |
| **9** | | | | | dPBS | 13 | 13 | 2/13 | 2/2 | 2/2 |
| **10** | | | | | Resuspension Buffer – 49°C | 7 | 7 | 0/7 | - | - |
| **11** | | | | | dPBS – 49°C | >500* | 20 | 0/20 | - | - |
| **12** | | | | *mnuA* mutant | Resuspension Buffer | 166 | 20 | 1/20 | 1/1 | 1/1 |
| **13** | | | | | dPBS | >300 | 20 | 0/20 | - | - |
| **14** | | | | | Resuspension Buffer – 49°C | 98 | 20 | 4/20 | 2/4 | 1/2 |
| **15** | | | | | dPBS – 49°C | >300* | 20 | 0/20 | - | - |
| **16** | - | - | - | Both | Resuspension Buffer | 0 | - | - | - | - |

[a] The number of yeast transformants analyzed by simplex PCR and multiplex PCR is reported, as well as the number of positive clones obtained.
[b]For control conditions (2 to 4) corresponding of plasmid (pMT85) transformation control protocol, the number of yeast transformants remained identical in addition of *M. agalactiae mnuA* mutant cells (>1000), but drastically decreased in addition of *M. agalactiae* WT cells.
*Lots of small colonies were observed as false transformants.

**Table S11: Mutations analysis with Galaxy after whole genome sequencing of genome tranplants.**
(Excel document).
